# *mKast* knockout honey bees show defects in brain functions underlying homing ability

**DOI:** 10.1101/2025.07.23.666469

**Authors:** Hiroki Kohno, Takeo Kubo

## Abstract

Honey bees visit food sources up to several kilometers away from their hives, which is underpinned by their sophisticated learning and memory, and cognitive abilities. However, the molecular and neural bases for these advanced brain functions remain obscure. Here, we focused on *mKast*, a gene preferentially expressed in the optic lobes, a visual center, and a specific Kenyon cell subtype in the mushroom bodies, a higher-order center, in the honey bee brain. We successfully produced homozygous mutant honey bee workers by crossing individuals mutated with CRISPR/Cas9 targeting *mKast*. Through behavioral analyses of *mKast* mutants using a new conditioning paradigm and a visual response assay, we found that *mKast* functions in bimodal learning and memory based on olfactory and visual information, and direction-specific motion sensing. We also found that *mKast* homozygous mutants have defects in survival outside the colony. These findings suggest that *mKast* modulate brain functions underlying homing ability that is essential for nidificating hymenopteran species.

## Introduction

Living in the nest is an important behavioral trait for many social animals and is considered a prerequisite for the acquisition of eusociality (*1–3*). The infraorder Aculeata (Insecta: Hymenoptera) is characterized with its nidification behaviors (*4*). The aculeate species includes all eusocial hymenopteran species, such as honey bees, ants, and hornets, and have an advanced learning and memory, and cognitive abilities to remember the location of their nests to return from foraging to raise their broods (*5–8*). The European honey bee (*Apis mellifera*) forms colonies and lives in hives, and its workers forage for nectar and pollen by memorizing the locations of their hives and food sources several kilometers away (*9*, *10*). The honey bee has long been studied as a model insect for the study of the molecular and neural basis of learning and memory, and social behaviors.

Mushroom bodies (MBs) are a higher-order brain center in insects and have been the focus of considerable attention in molecular ethological studies of honey bees (*11–14*). MBs are elaborated in Aculeata among hymenopteran insects, and the honey bee MBs consist of approximately 300,000 cells among approximately one million cells in the brain (*15*, *16*). The honey bee MBs change the volume of the neuropil and gene expression profiles associated with the age-dependent division of labor of workers, from nursing in the hive (nurse bees) to foraging outside the hive (foragers), suggesting that they play roles in regulating social behaviors (*12*). Honey bee MBs comprise intrinsic neurons termed Kenyon cells (KCs), which are further classified into three subtypes of Class I KCs (large-, middle-, and small-type KCs [lKCs, mKCs, and sKCs]), which differ in size and location of their somata and gene expression profile, and Class II KCs (*17*). The number of KC subtypes has likely increased with behavioral evolution in hymenopteran insects, from one in the phytophagous suborder Symphyta, two in the parasitic Parasitica, and three in Aculeata (*16*). This implies that the acquisition of three KC subtypes is associated with the evolution of behavioral traits specific to aculeate species.

So far, lKCs are suggested to be related to learning and memory ability, because genes involved in Ca^2+^-signaling, which underlies the learning and memory, are preferentially expressed in lKCs, and RNAi against *Ca^2+^/calmodulin-dependent protein kinase II* (*CaMKII*), which is also preferentially expressed in the honey bee lKCs, impairs long-term memory in the honey bee (*18*). In addition, neural activity mapping using an immediate early gene, *kakusei*, revealed that sKCs and some mKCs were activated during the foraging flight, suggesting that these KC subtypes are related to the brain functions required for foraging behavior (*19*, *20*). *Ecdysone receptor* is preferentially expressed in sKCs in the honey bee brain and is induced by foraging flight possibly to repress lipid metabolism through the transcriptional regulation of downstream genes (*21*, *22*). However, the function of genes preferentially expressed in each KC subtype and thus the function of each KCs, especially that of mKCs, remain elusive, mainly because of the lack of effective gene knock out techniques in honey bees until recently (*23*).

Here, we focused on *middle-type Kenyon cell-preferential arrestin-related protein* (*mKast*), which is preferentially expressed in the mKCs of honey bee MBs. Although *mKast* was originally identified during a search for genes that are highly expressed in the honey bee optic lobes (OLs), a visual center of the insect brain, a detailed analysis of its expression pattern in MBs led to the discovery of mKCs (*19*). *mKast* encodes a protein that have an arrestin domain and belongs to the α-arrestin family that plays multiple functions, including downregulation of activated membrane proteins (*24*, *25*). However, its amino acid sequence is not well conserved, including in hymenopteran insects, and its function is unknown in insects. *mKast* that is selectively expressed in the brain among adult body parts, and in the adult brain, mainly in OLs and mKCs (Fig. 1A) (*19*, *26*). *mKast* begins to be expressed in the brain from late pupal stages (P7) and thus functions specifically in the adult brain (*26*). These expression patterns imply that *mKast* is involved in the behavioral regulation of adult worker honey bees.

**Fig. 1.**
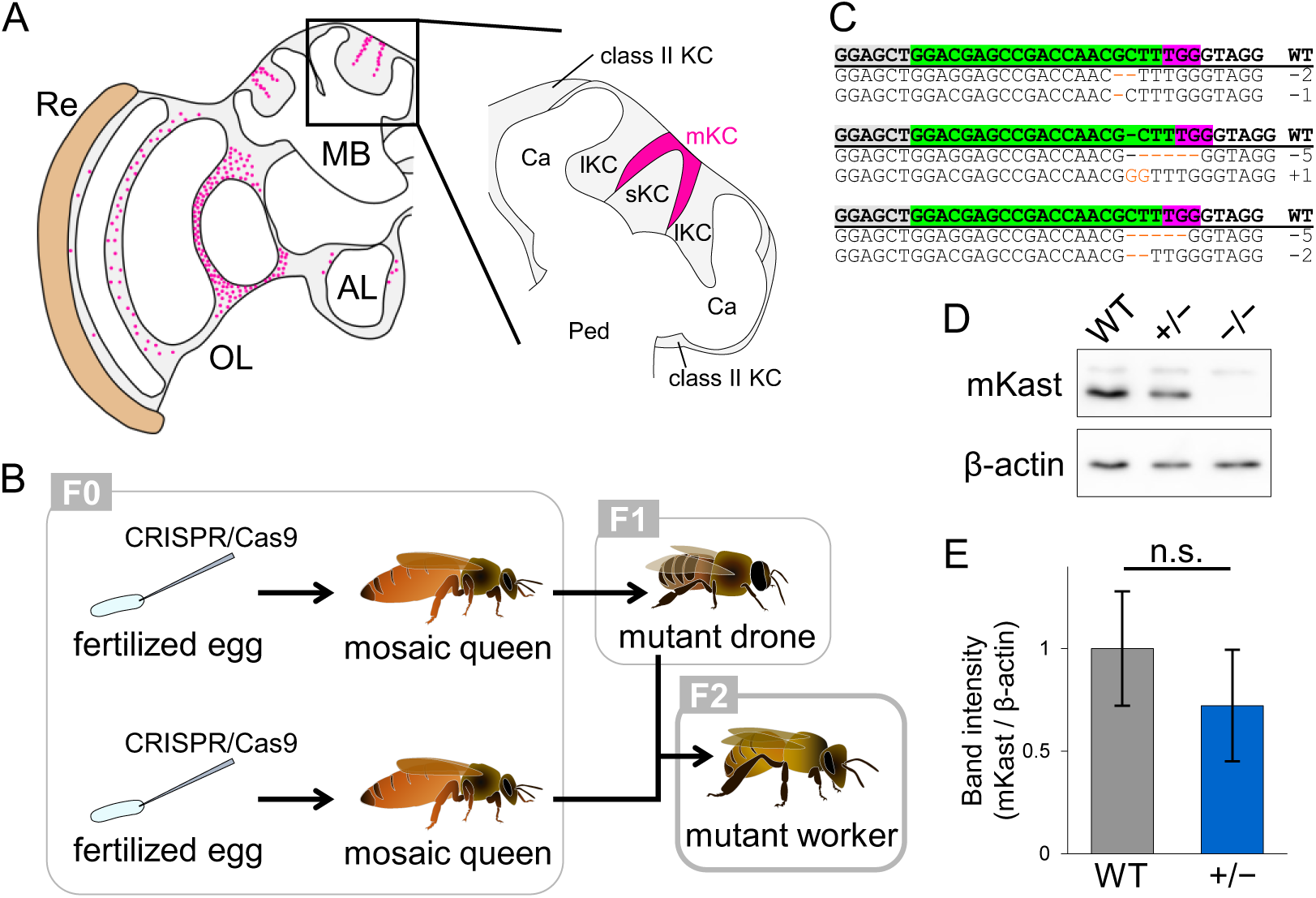
Production of *mKast* homozygous mutant workers. (**A**) Distribution of *mKast*-expressing cells (magenta dots and magenta region) in the adult brain hemisphere (left) and in the MB (right). MB, mushroom body, AL, antennal lobe, OL, optic lobe, Ce, compound eye, Ca, calyx, Ped, pedunculus. (**B**) Process to produce homozygous mutant workers (F2) by CRISPR/Cas9. (**C**) Indels detected at sgRNA target site in homozygous mutant workers. Letters marked in green, magenta, and gray in WT sequence indicate sgRNA target site, PAM sequence, and exon of *mKast*, respectively. Orange letters and dashes indicate the inserted and deleted nucleotide sequences, respectively. (**D**) Results of immunoblot analysis using brain homogenates of WT, hetero-(+/−) and homozygous (−/−) *mKast* mutants and anti-mKast (top) and anti-β-actin (bottom) antibodies. (**E**) The intensity of the bands for mKast protein relative to those of β-actin in immunoblot analysis (D) in WT and heterozygous mutants. n = 3 for each genotype. n.s., not significant by t-test.

Here, we successfully produced homozygous mutant workers by crossing individuals whose genomes were edited by CRISPR/Cas9 targeting *mKast*. The function of *mKast* was inferred from the localization of the mKast protein in the MBs and OLs, and by comparative analysis of honey bee and *Drosophila* OL single-cell (sc) RNA-seq data. Using *mKast* mutant workers and behavioral assays, including a newly established paradigm, we found that *mKast* functions in bimodal learning and memory based on olfactory and visual information, direction-specific motion sensing, and survival outside the colony. Our findings suggested that *mKast* modulates brain function which are presumed to underlie the homing ability of honey bees.

## Results

### Production of *mKast* homozygous mutant workers

Some previous studies have analyzed the phenotypes of honey bee workers (F0) artificially reared from fertilized eggs mutated with CRISPR/Cas9 with high efficiency (*23*, *27*). However, this method has some disadvantages; it is unclear whether mutations are introduced into all the cells of the entire body of each individual used for phenotypic analysis, and artificially reared workers differ from workers reared in normal colonies in terms of gene expression, physiological state, and behavior (*28*, *29*). Therefore, we intended to produce *mKast* homozygous mutant workers by crossing genome-edited individuals produced using CRISPR/Cas9 (Fig. 1B and Fig. S1, see Materials and Methods). Mosaic queens (F0) were produced from fertilized eggs injected with Cas9 protein and sgRNA targeting *mKast*. Some mosaic queens (F0) with high mutation efficiency were selected by genotyping (Fig. S1), and were forced to lay unfertilized eggs that develop into drones (male honey bees) by CO_2_ treatment. Semen collected from sexually mature drones (F1) was used for artificial insemination of mosaic queens (F0) that were produced separately. Hetero- and homozygous mutant workers (F2) were produced from the artificially inseminated mosaic queens (F0).

Sequencing analysis of the PCR products of the homozygous mutants revealed indels, resulting in frameshift mutations in the sgRNA target sites of both alleles (Fig. 1C). Immunoblotting using crude brain protein extracts and anti-mKast antibodies showed that the band for mKast protein was completely absent in the homozygous mutant (Fig. 1D). Although bands for mKast were detected in the heterozygous mutants, their band intensities were approximately 70% of those in the WT (Fig. 1E). These results showed that *mKast* was knocked out in the homozygous mutant workers. We sequenced the genomic regions of potential off-target candidates selected using Cas-OFFinder (*30*) (Table S1). A single nucleotide mismatch in the PAM sequence was detected in one allele of a candidate sequence (Candidate #2 in Table S1) in some hetero- and homozygous mutants. Although we cannot exclude the possibility that this mutation was caused by an off-target double-strand break (DSB), it is more likely that the detected mismatch in the PAM sequence is due to a single nucleotide polymorphism. This is because DSB by CRISPR/Cas9 and subsequent indel mutations usually occur three bases upstream of the PAM sequence (*31*). Even if this mismatch was caused by off-target DSB, it is unlikely that this mutation would affect the results of future analyses because the candidate sequence is located in an intron of a gene. Genomic regions corresponding to other off-target candidates were identical to the reference genome in all individuals (Table S1). This indicates that there were no detectable off-target mutations in the *mKast* mutants produced in this study.

To investigate the effect of *mKast* KO on the gene expression profile of the brain, we performed RNA-seq analysis of the MBs and OLs of WT and mutant workers. In both MBs and OLs, gene expression profiles were more similar between heterozygous and homozygous mutants than between the mutants and the WT (Fig. S2A, B). The most DEGs were detected between the WT and mutants in both MBs and OLs, and relatively few DEGs were detected between the hetero- and homozygous mutants (Fig. S2A, B, Tables S2–S5). Gene ontology (GO) enrichment analysis showed that many GO terms were commonly detected in the DEGs of MBs and OLs, most of which were essential for cellular functions (Fig. S2C). GO terms specific to the MB and OL were also detected, suggesting that *mKast* has both brain-region-specific and common functions. In contrast, GO analysis of the DEGs of MBs and OLs between heterozygous and homozygous mutants showed that only some GO terms were commonly detected in MBs and OLs (Fig. S2D, E). Meanwhile, most GO terms were not common between DEGs of MBs and OLs. DEGs of MBs included genes related to behaviors, cell morphology, and various signaling and metabolism, whereas DEGs of OL were enriched with genes related to membrane transport, such as transmembrane transport and Golgi organization. These results are consistent with the fact that mKast belongs to the α-arrestin family, which not only regulates activities of membrane receptors such as G-protein-coupled receptors (GPCRs) but also interacts with diverse proteins (*24*, *32*, *33*), and suggest that mKast interacts with different factors depending on brain regions.

### Localization of mKast protein in the WT and mutant brains

We previously reported the localization of mKast protein in the worker honey bee brain by immunohistochemistry (IHC) (*26*), however, antibody-specific signals were detected in the entire region inside the MB calyces, which was inconsistent with the mKC-preferential expression pattern of *mKast* by *in situ* hybridization (ISH) (*19*). In addition, the distribution of mKast protein in the MB calyces and pedunculus was not observed because of the relatively high background. Therefore, we investigated the mKast IHC signals that are detectable in WT but not in homozygous mutant brains, which represent genuine mKast protein localization (Fig. 2 and Fig. S3).

**Fig. 2.**
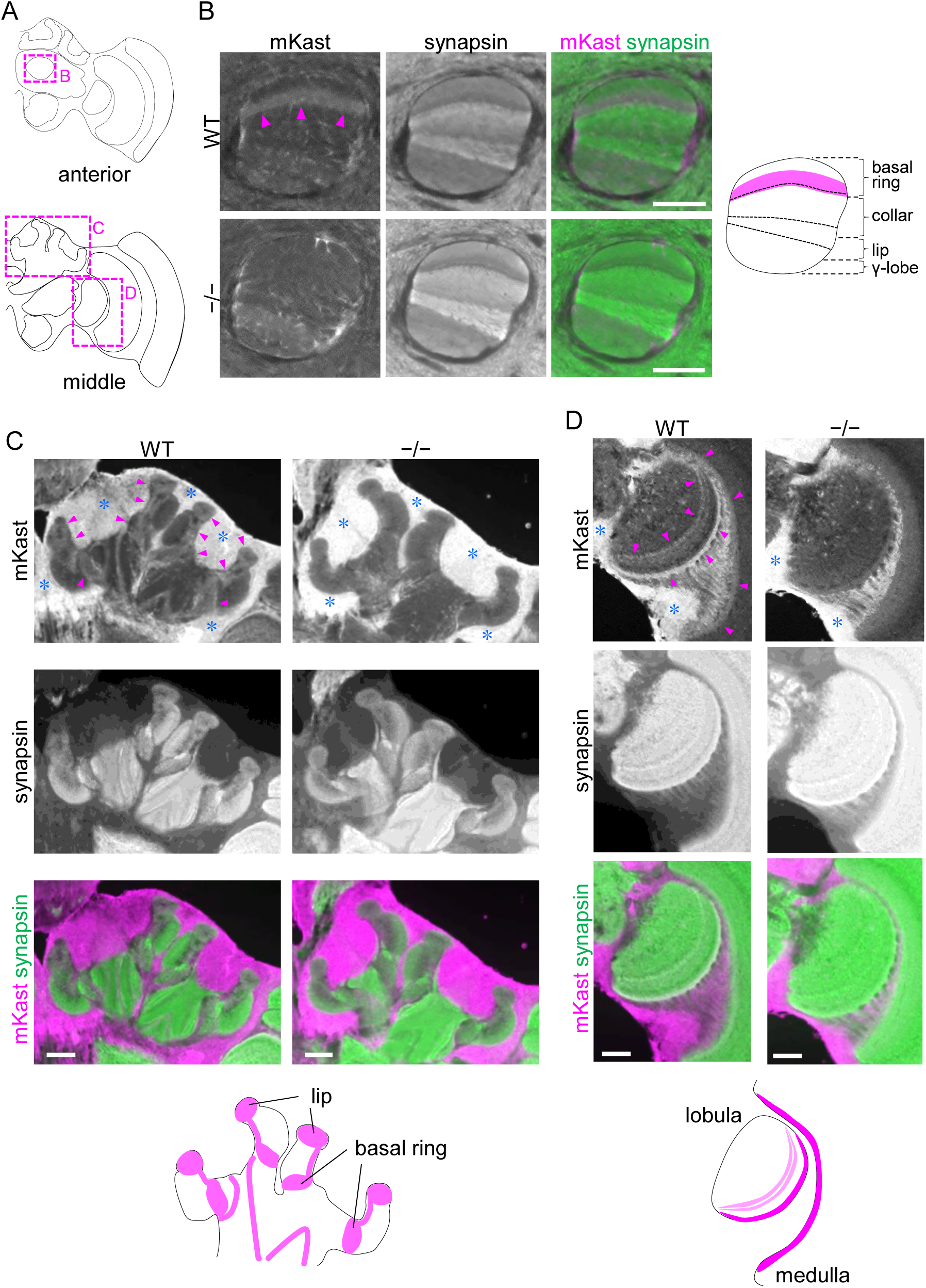
Localization of mKast protein in adult worker brain. (**A**) Schematic diagrams of anterior (top) and middle (bottom) regions in the adult honey bee brain. Magenta dotted boxes indicate areas corresponding to panels B–D. (**B–D**) Results of IHC of WT and homozygous mutant workers. mKast and synapsin IHC signals and their merged images in the MB vertical lobe (B), MB calyces (C), and OL (D) are shown. Magenta arrowheads indicate the mKast IHC signals specific to WT, and blue asterisks indicate non-specific fluorescent signals in the cell body region. Magenta in schematic diagrams on the right (B) and at the bottom (C and D) shows the localization of mKast protein in each brain region. The dotted line in the schematic diagram of the vertical lobe (B) represents the layers corresponding to the calyceal compartments. Scale bar, 100 µm.

In the vertical lobes, the output region of the MBs, of WT and heterozygous mutants, IHC signals were detected in the dorsal layer (Fig. 2B and Figs. S3A to C, S4A, B). In honey bee MBs, KCs that extend their dendrites into each calyceal compartment (the lip, collar, and basal ring) extend their axons into distinct layers of the vertical lobe (Fig. S4C) (*34*): KCs extending their dendrites into basal rings project their axons to the most dorsal layer of the vertical lobe, while those extending their dendrites into collars project their axons to a layer ventral to that of basal rings. Similarly, KCs extending their dendrites into lips project their axons to a layer ventral to that of collars. According to the IHC signals of FMRFamide, which has been reported to correspond precisely to the vertical lobe layer and the calyceal compartment, we confirmed that mKast IHC signals were detected in the layer which corresponds to a part of the region where KCs that extend their dendrites into the basal rings extend their axons (Fig. S4D, E) (*34*). In the MB calyces, the input region of the MBs, signals were detected not only in the basal rings but also in the lips, the latter of which possibly representing the localization of mKast at the presynapse of medial antenno-protocerebral tract, one of the olfactory pathways that connect the antennal lobes (ALs), an olfactory center of the insect brain, and MBs (Fig. 2C and Fig. S3D to F) (*26*, *35*). In the OLs, signals were detected in the innermost layer of the medulla and three layers of the lobula (Fig. 2D and Fig. S3G to I). These signals in the MBs and OLs were absent in homozygous mutants. Strong fluorescence in areas where cell bodies were present was detected even in the homozygous mutants (Fig. 2, blue asterisks), indicating that they did not represent mKast protein localization. Therefore, the discrepancy between mKast IHC and ISH inside the MB calyces in previous studies is due to this false-positive IHC signal. These results suggest that *mKast*-expressing neurons in the MBs, mKCs, extend their dendrites to the basal rings that receive sensory inputs from both ALs and OLs (*34*), and that those in the OLs are related to the function of the lobula and medulla, such as detection of movement, color, and shape (*36*).

### Function of *mKast* in learning and memory abilities

Because MBs are involved in learning and memory in various insects including honey bees (*13*, *37*), we considered the possibility that *mKast* is involved in learning and memory based on multimodal sensory information, which enables bees to forage efficiently by learning multimodal floral stimuli (*38*, *39*). Therefore, we investigated the defects of *mKast* mutants in associative learning of a combination of olfactory and visual information.

First, we created an experimental setup to present four combinations of colors (blue or green) and odors (linalool or 2-octanol), as conditioned stimuli (CSs), and 50% sucrose or 1M NaCl solution as unconditioned stimuli of reward (US+) or punishment (US−), respectively (Fig. 3A and Fig. S5). This setup enabled us to present only CSs, and combinations of CSs and USs (US+ at the area with a combination of green and linalool, and US− at the other three areas with other CSs) to a bee in the arena (Fig. 3B and Fig. S5, see Materials and Methods). One training trial consisted of the presentation of CSs for 5 s, followed by the presentation of combinations of CSs and USs until the bee in the arena reached the US+. This trial was repeated eight times at 1-min intervals (Fig. S5F, Movie S1). A memory test was conducted approximately 24 h after the end of training, in which only the CSs were presented for 15 s (Fig. 3B, Movie S2). In addition to this paired group, an unpaired group was prepared in which CSs and USs were presented separately during training (see Materials and Methods). We calculated the preference index (PI) for the area with the CS conditioned with US+ (CS+: green and linalool) during the presentation of CSs in each trial in the training and in the memory test. The paired group tended to show a higher PI for the CS+ area than the unpaired group during training. However, no significant difference was observed between them because the PI varied largely among individuals (Fig. 3C). In contrast, in the memory test, the PI for the CS+ area was significantly higher in the paired group than in the unpaired group (Fig. 3D). The CS+ area was not presented clockwise in the memory test, indicating that the bees did not learn the rotation rule during the training, where the CS+ area was rotated 90° clockwise with each trial (Fig. S5F). This suggested that the paired group learned the correct combination of color and odor.

**Fig. 3.**
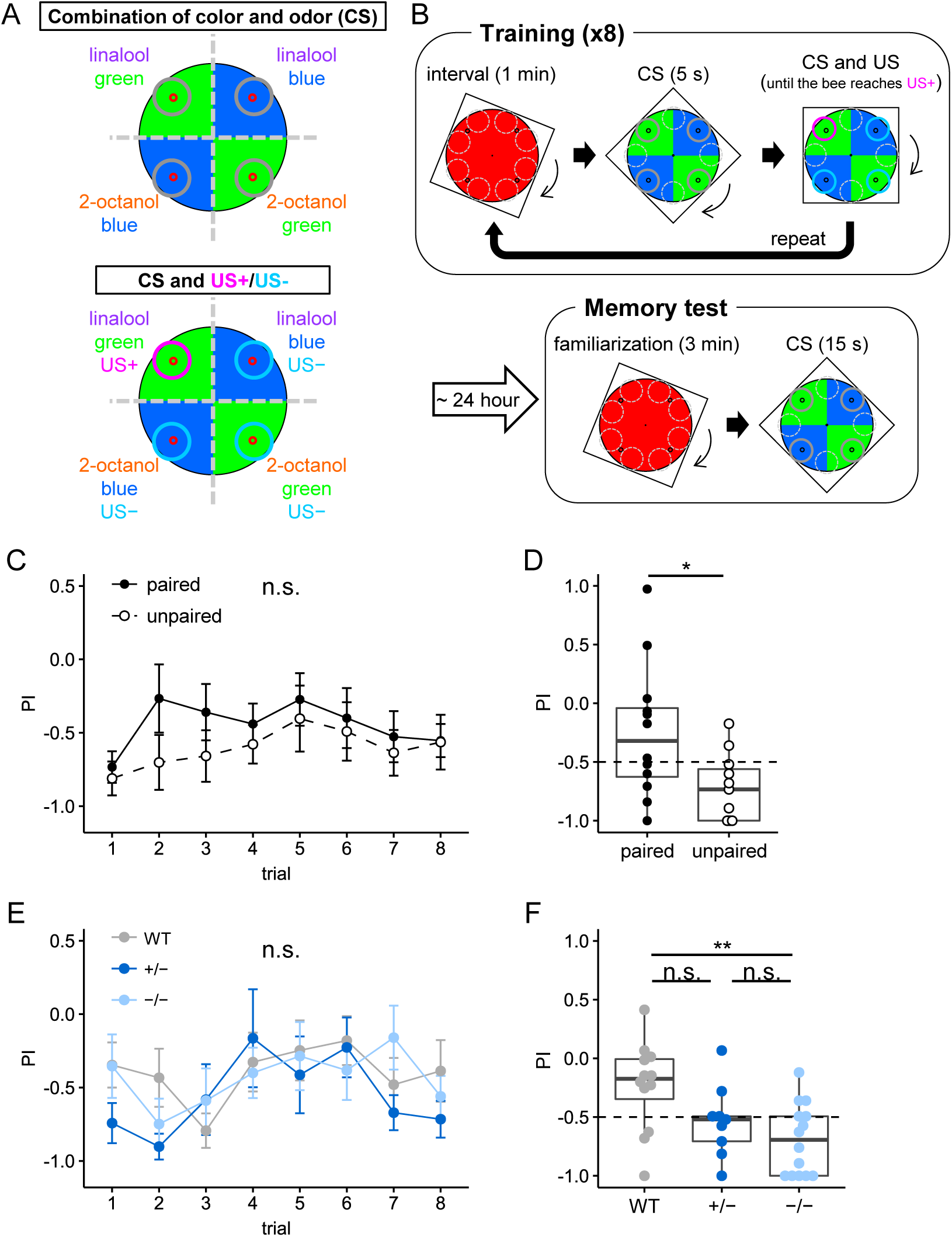
Effect of *mKast* knockout on bimodal conditioning analyzed using a newly established experimental system. (**A**) Schematic diagrams of the presentation of CSs (top) and combinations of CSs and USs (US+: 50% sucrose, US−: 1M NaCl) (bottom) in the arena. The odors and US are presented to a bee in the arena through holes at the bottom of the arena (red circles) from dishes (gray, magenta, and light blue circles) placed between the arena and the monitor. (**B**) Schemes of the bimodal conditioning experiment of paired group. (**C, D**) PIs of the bees in paired and unpaired groups during the presentation of CS in each trial in the training (C) and in the memory test (D). n = 12 and 11 in the paired and unpaired groups, respectively. Mann–Whitney U test (C), and Welch’s t-test (D), * p < 0.05, n.s., not significant. (**E, F**) PIs of bees with each genotype during CS presentation in each trial in the training (E) and the memory test (F). n = 12, 9, and 14 in WT, hetero- and homozygous mutants, respectively. Kruskal–Wallis test (E), and Tukey HSD test (F), ** p < 0.01, n.s., not significant. The dotted line (D, F) shows the expected value of PI when a bee moves randomly (PI = −0.5).

Using this experimental system, the learning and memory abilities of color and odor combinations in the WT and mutants were examined. During the training, the PI for the CS+ area was not significantly different between the genotypes (Fig. 3E). The time to reach US+ in each trial decreased as training progressed for all genotypes (Fig. S6A). In contrast, in the memory test, homozygous mutants showed a significantly reduced PI for the CS+ area compared to the WT (Fig. 3F). The PI for the CS+ area of heterozygous mutants was not significantly different from that of either the WT or homozygous mutants. The distance that the bees moved in the arena during the memory test did not differ between genotypes (Fig. S6B), because bees that had not learned the CS+ continued to walk around the arena, whereas learned bees also continued to search for US+ in the arena, mainly in the CS+ area. This suggested that the low PI for the CS+ area in homozygous mutants was not due to decreased mobility. In addition, we conducted olfactory or visual proboscis extension reflex (PER) conditioning assay (*40*, *41*) in WT and mutants using the same odors (linalool and 2-octanol) or colors (blue and green) used for bimodal conditioning as the CS to examine whether the mutant workers also exhibit defects in unimodal learning and memory abilities. However, the homozygous mutant had a comparable level of learning and memory ability as the WT in both olfactory PER associative learning and visual PER associative learning (Fig. S7). Therefore, although homozygous *mKast* mutants can learn and memorize unimodal sensory (olfactory or visual) information and can recognize odors and colors presented in bimodal conditioning assays, they have defects in the learning and memory of bimodal sensory information.

### Identification of *mKast*-expressing OL neuron by comparative scRNA-seq analysis

Because our knowledge of the function of OL neurons in honey bees is limited, we aimed to identify *mKast*-expressing OL neurons and infer their functions by comparing honey bee OL scRNA-seq data with those of *Drosophila*, in which the classification and function of OL neurons have been studied in detail (Fig. 4A).

**Fig. 4.**
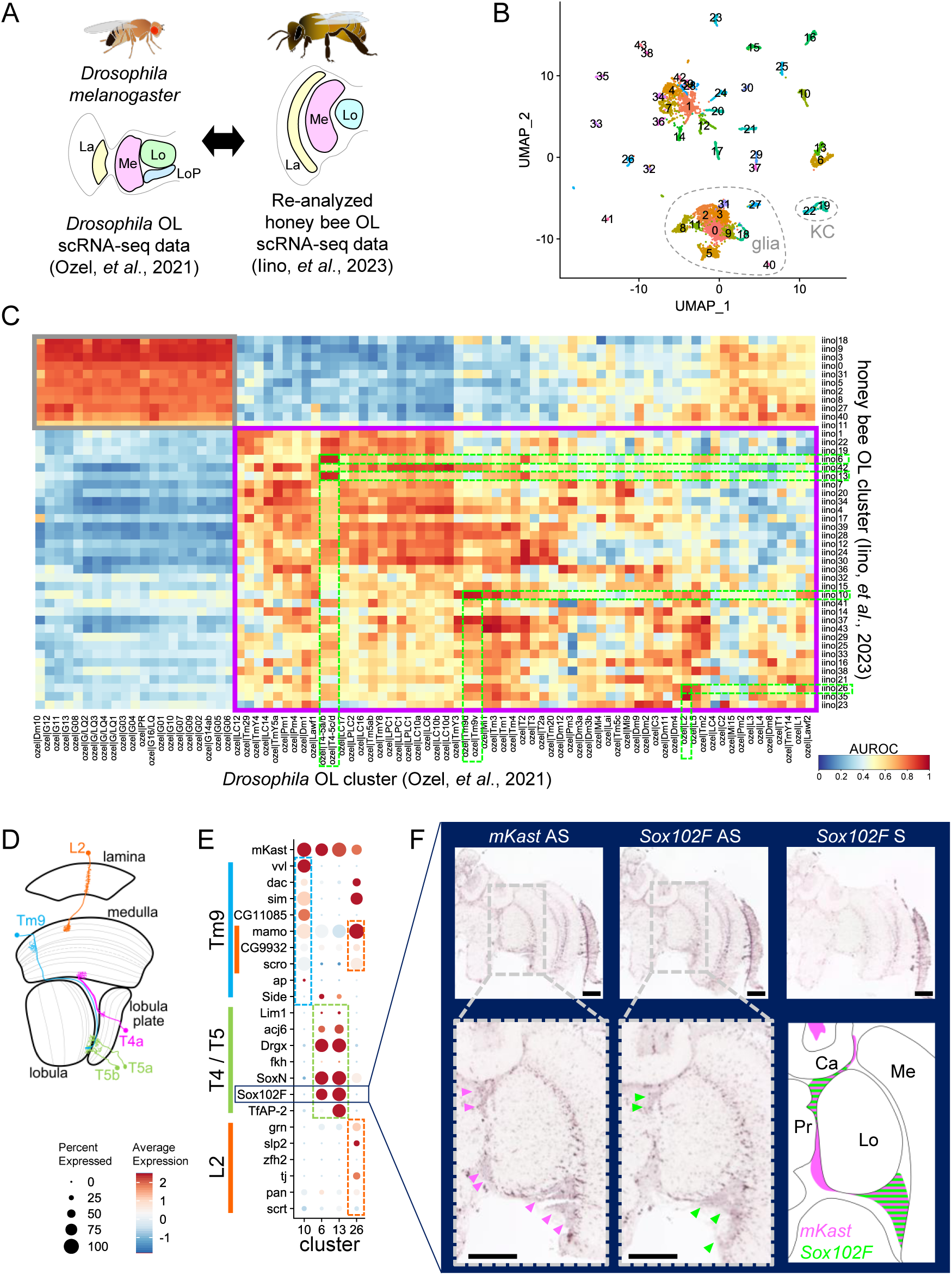
Identification of *mKast*-expressing OL neurons by comparative analysis of OL scRNA-seq data between the honey bee and fruit fly. (**A**) Scheme for data analysis. La, lamina, Me, medulla, Lo, lobula, LoP, lobula plate. (**B**) UMAP visualization of clusters of re-analyzed honey bee OL scRNA-seq data. (**C**) Heatmap of AUROC scores between *Drosophila* and honey bee OL clusters calculated using MetaNeighbor. Gray and magenta boxes indicate glia and neurons, respectively, and the yellow-green dotted lines indicate the *mKast*-expressing clusters in the honey bee and *Drosophila* clusters that showed high AUROC scores with these clusters. AUROC, area under the receiver operating characteristic curve. (**D**) Schematic diagram of the *Drosophila* OL neurons that showed high AUROC score with the honey bee *mKast*-expressing OL cells in the *Drosophila* brain. (**E**) A dot plot showing the expression levels of marker genes for *Drosophila* L2, Tm9, and T4/5 neurons in the honey bee *mKast*-expressing OL cell clusters. Light blue, yellow-green, and orange bars indicate marker genes in Tm9, T4/5, and L2 neurons, respectively. (**F**) ISH of *Sox102F* and *mKast* using serial honey bee brain sections. Magenta and yellow-green arrowheads indicate signals for *mKast* and *Sox102F*, respectively. The lower right panel schematically shows the expression patterns of *mKast* (magenta) and *Sox102F* (yellow-green). AS, antisense probe, S, sense probe, Ca, calyx, Me, medulla, Lo, lobula, Pr, protocerebrum. Scale bar, 200 µm.

We reanalyzed the scRNA-seq data of honey bee OLs from a previous study (*22*) and clustered the cells into 44 clusters (Fig. 4B). Based on the expression patterns of marker genes for neurons, glia, and KCs (*42*, *43*), 33 clusters were annotated as neurons, including two KC clusters that may have been contaminated during dissection and 11 clusters as glia (Fig. S8A). By calculating the AUROC scores between these honey bee OL clusters and those in the *Drosophila* OL scRNA-seq data (*44*) using MetaNeighbor (*45*), we searched for *Drosophila* clusters to which each *mKast*-expressing OL neuron cluster in the honey bee scRNA-seq data showed high similarities. The *mKast*-expressing honey bee OL clusters (Fig. S8B) showed the highest AUROC scores for *Drosophila* Tm9d, T4-5a/b, and L2 OL neurons (Fig. 4C, D and Fig. S8C, Tables S6, S7). Honey bee OL cluster 10, which had the highest AUROC score with Tm9d, and honey bee OL clusters 6 and 13, which had the highest AUROC score with T4-5a/b, also showed high, almost equal to the highest, AUROC scores with Tm9v and T4-5c/d, respectively (Fig. 4C, Table S7). The marker genes of these cell types in *Drosophila* (*44*) were selectively expressed in the *mKast*-expressing clusters among the honey bee OL clusters (Fig. 4E and Fig. S8D). The ISH of a marker gene of T4/5 cells, *Sox102F*, and *mKast* in serial brain sections showed that areas in which *mKast*-expressing neurons exist overlapped with those of *Sox102F*-expressing neurons (Fig. 4F), supporting cell type estimation by MetaNeighbor. In addition, T4 neurons connect the innermost layer of the medulla to layers in the lobula plate, and T5 neurons connect layers in the lobula plate to those in the lobula (Fig. 4D), which are consistent with the layered mKast IHC signal patterns in the lobula and medulla (Fig. 2D). These results suggested that the *mKast*-expressing cells in honey bee OLs were L2, Tm9, and T4/5 cells. *Drosophila* L2, Tm9, and T4/5 cells are involved in motion sensing and the four subtypes (a, b, c, and d) of T4/5 cells respond selectively to four different directions: upward, downward, forward, and backward (*46*, *47*). Therefore, it is plausible that *mKast*-expressing OL neurons are involved in motion sensing in honey bees.

### Function of *mKast* in motion sensing

Honey bees show an antennal response (AR) to motion stimuli on the compound eye by tilting their antennae in the opposite direction to the motion to perceive sensory information in the direction they are moving (*48*). Therefore, we examined the defects of *mKast* homozygous mutants in the motion sensing using the experimental system developed in our previous study (*49*). We quantified the ARs in response to motion stimuli in the transverse and coronal planes in the WT and mutants by measuring the angles of the antennae from the head midline (Fig. 5A to D and Fig. S9A, B). When vertical motion stimuli were presented, the antennal angles in the transverse plane tended to be smaller for upward motion at approximately 60° from the midline. Meanwhile, they tended to be larger for downward motion at approximately 95° and intermediate during spontaneous movement at approximately 85° in all genotypes (Fig. 5A, C and Fig. S9A, Movie S3). However, the angle during spontaneous movement was significantly larger in the homozygous mutants than in the WT (Fig. 5C), and the angle for downward motion was not significantly different from that during spontaneous movement in the homozygous mutants (Fig. S9A). When horizontal motion stimuli were presented, the antennal angles in the coronal plane tended to be smaller for backward motion at approximately 40°, while they tended to be larger for forward motion at approximately 70°, and intermediate during spontaneous movement at approximately 60° in all genotypes (Fig. 5B, D and Fig. S9B, Movie S4). However, the antennal angle was significantly smaller in the homozygous mutants than in the WT during the presentation of the backward motion stimulus (Fig. 5D).

**Fig. 5.**
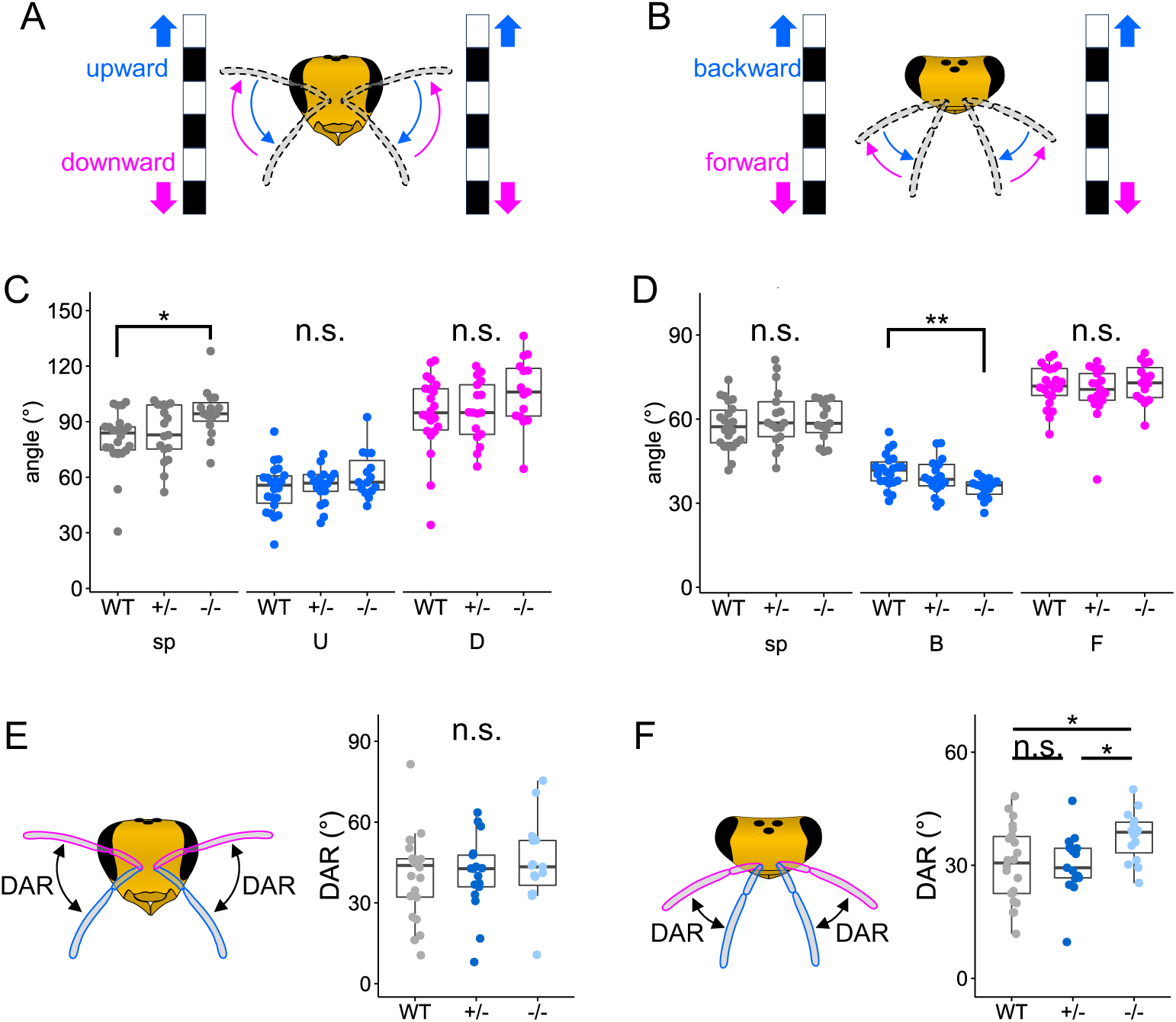
Effect of *mKast* knockout on ARs to motion stimuli. (**A, B**) Schematic diagrams of the presentation of motion stimuli (moving latices) to a fixed bee in the transverse plane (A) and in the coronal plane (B). (**C, D**) Comparison of antennal angles during the presentation of motion stimuli and spontaneous movement in the transverse plane (C) and coronal plane (D) between genotypes. Tukey HSD test, ** p < 0.01, * p < 0.05, n.s., not significant. (**E, F**) Comparison of DAR in the transverse plane (E) and coronal plane (F) between genotypes. Tukey HSD test, * p < 0.05, n.s., not significant. n = 22, 17, and 15 for WT, heterozygous mutants, and homozygous mutants, respectively.

Consistent with these results, the angle for downward motion did not differ significantly from that during spontaneous movement in the homozygous mutants (Fig. S9C, D), and the direction-specific antennal response (DAR), which is the difference in the angles of the antennae for upward and downward motion stimuli or backward and forward motion stimuli, was significantly higher in the homozygous mutants than in the heterozygous mutants or the WT in the coronal plane (Fig. 5E, F). In contrast, no significant differences were observed in the maximum/minimum angles and angular range of the antennae during spontaneous movement in the transverse and coronal planes among the genotypes (Fig. S9E to J). This suggests that the differences in ARs detected between the genotypes were not due to defects in antennal mobility. Therefore, homozygous mutants have defects in ARs to motion stimuli in a direction-specific manner, which is consistent with the notion that *mKast*-expressing OL neurons are involved in motion sensing.

### Effects of *mKast* KO on survival rate

Finally, we compared the survival rates of the WT and mutant workers. When newly emerged WT and mutant workers were reared in closed plastic cages mimicking the in-hive conditions with free access to honey, water and pollen substitutes (Fig. 6A), no differences were observed in the survival rates between the genotypes (Fig. 6B). This suggests that *mKast* is dispensable for normal lives in the colony, which is consistent with our previous report that *mKast* is dispensable for normal development. In contrast, when newly emerged mutants were introduced into a normal colony set in the flight room (Fig. 6C), although the proportions of hetero- and homozygous mutants were the same immediately after emergence, the proportion of homozygous mutants was significantly reduced compared to that of heterozygous mutants when individuals inside and outside the hive were collected and genotyped more than 14 d after emergence (Fig. 6D). Because bees can freely enter and leave the hive when reared in the colony, homozygous mutants may have some defects in survival outside the hive and/or in returning to the hive. Therefore, *mKast* may be involved in the regulation of behavior and physiological state outside the colony.

**Fig. 6.**
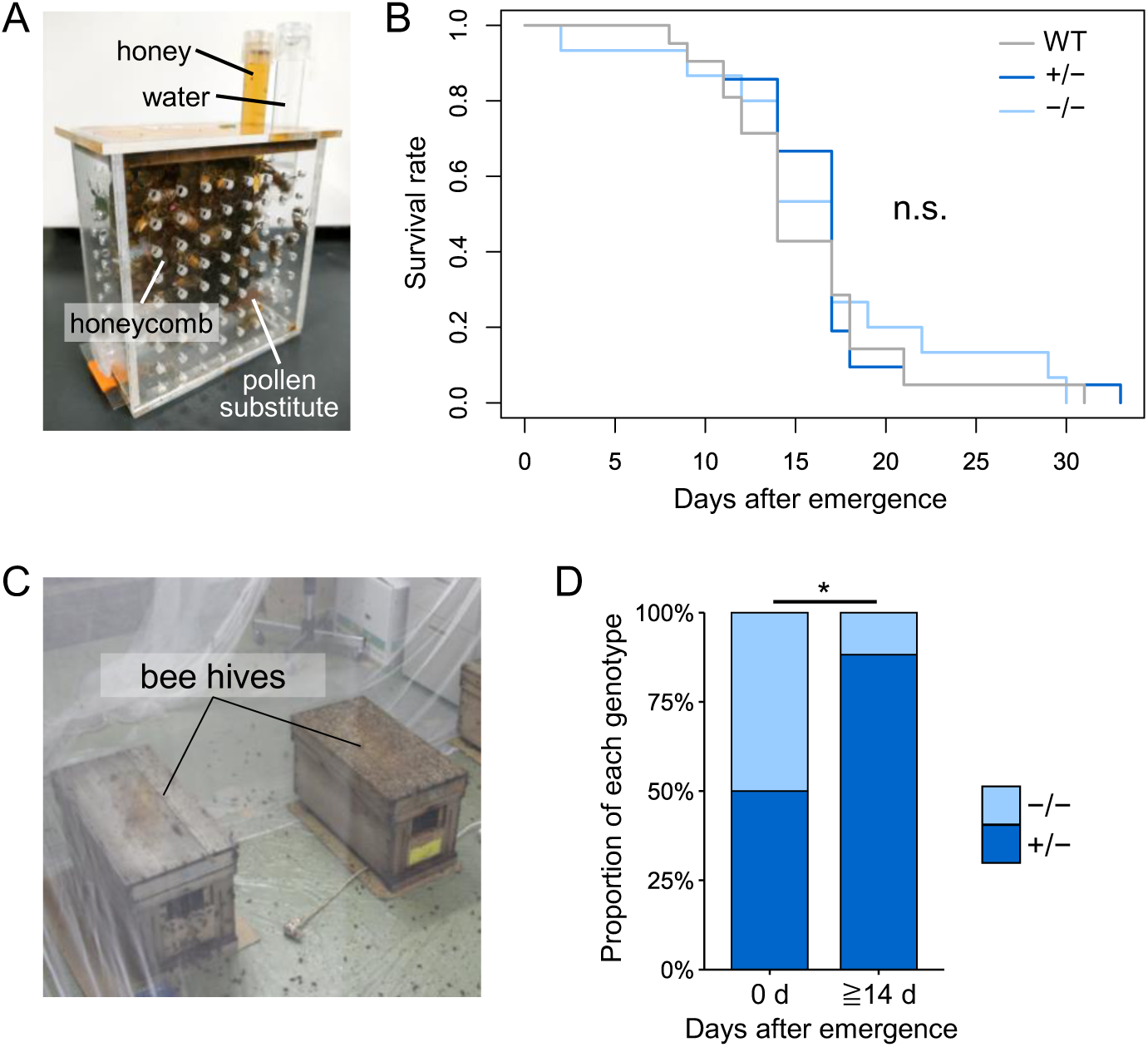
Survival rates of WT and mutants under different rearing conditions. **(A)** plastic cage containing a honeycomb. Honey, water, and pollen substitute are supplied *ad libitum*. **(B)** Survival curves for each genotype reared in plastic cages. n = 21, 21, and 15 for WT, heterozygous mutants, and homozygous mutants, respectively. Log Rank test, n.s., not significant. (**C**) Hives set inside the flight room. (**D**) Percentage of heterozygous and homozygous mutants immediately after emergence (0 day) or reared 14 days or more after the emergence in colonies set inside the flight room. n = 32 and 17 for 0 day and 14 days or more after emergence, respectively. Fisher’s exact test, * p < 0.05.

## Discussion

In the present study, we focused on *mKast*, a gene of unknown function that is selectively expressed in the adult worker brain and revealed its function in behavioral regulation of honey bees using homozygous mutant workers produced by crossing genome-edited individuals. Since *mKast* is mainly expressed in the OLs and the MBs in the adult brain, it is plausible that *mKast* is involved in the learning and memory of multiple sensory information in mKCs in the MBs, the detection of movement in a specific direction in particular OL neurons (e.g., L2, Tm9, T4/5 cells), and the regulation of behavior and physiological states outside the colony through its function in MBs, OLs, or other cells expressing it.

The α-arrestin family, to which *mKast* belongs, has been shown to be involved in the regulation of the activity of membrane proteins such as GPCRs (*24*, *32*). This occurs by internalizing activated membrane receptors by endocytosis or by recruiting E3 ubiquitin ligase to direct the degradation pathway (*25*, *50*). Recent proteome analyses constructed protein–protein interactions network for α-arrestins comprised of hundreds of proteins and suggested their previously unknown molecular functions. In insect MBs, projection neurons from the ALs, which convey odor information, and dopaminergic and octopaminergic neurons, which convey information about the US, such as reward and punishment, provide inputs and regulate synaptic plasticity between KCs and MB output neurons (*51–53*). This results in the formation of associative memory for the odor. *mKast* may be involved in the learning and memory of multiple sensory information through the regulation of protein activities involved in synaptic plasticity, such as receptors for above-mentioned neurotransmitters, in mKCs, which extend their dendrites into basal rings where both olfactory and visual sensory information input.

The antennal angle during spontaneous movements in the transverse plane significantly increased in *mKast* homozygous mutants when compared with WT and thus did not differ from the angle when presented with a downward motion stimulus (Fig. 5C and Fig. S9A, C). The AR to the downward motion stimulus did not differ between WT and mutants, suggesting that *mKast* homozygous mutants could perceive and respond to downward motion stimuli. Therefore, this abnormal antennal angle during spontaneous movement in the homozygous mutants can be explained if *mKast* is involved in signaling that inhibits or shuts down information about downward motion stimuli. The homozygous mutants also showed excessive AR to the backward motion stimulus and thus had a larger DAR in the coronal plane (Fig. 5D, F). In *Drosophila*, T4/5 neuron subtypes (a–d) change their responses depending on the motion velocity (*47*). Therefore, it is possible that *mKast* functions in velocity dependency in T4/5a neurons which respond to backward motion stimuli (*47*), resulting in excess AR to a backward motion in the homozygous mutants. Another possibility is that *mKast* functions in L2 and Tm9, which are upstream of T4/5 neurons (*46*), lead to the phenotype observed in the homozygous mutants. The DAR in the transverse plane is increased by octopamine administration and decreased by serotonin administration in the lobula (*54*). The IHC signals for serotonin and octopamine exist in the lobula (*55–57*), consistent with the mKast IHC signals observed in this study. Therefore, the defects of *mKast* homozygous mutants in ARs can be explained if *mKast* is involved in the regulation of molecular cascades, leading to the exocytosis of these neurotransmitters. As the motion presented to the compound eye is opposite to the direction in which bees move under natural conditions: bees perceive downward motion stimuli when moving upward and backward motion stimuli when moving forward. Abnormalities in the perception of these optic flows hinder the ability of *mKast* homozygous mutants to perform proper foraging flights.

In the present study, no significant differences were observed between homozygous and heterozygous mutants in the bimodal conditioning assay and in part of the AR analysis, in which heterozygous mutants showed an intermediate phenotype between the WT and homozygous mutants (Figs. 3F, 5C, D). *mKast* protein was still expressed and localized in the brains of heterozygous mutants as well as in the WT (Fig. 1D and Fig. S3). However, although no significantly different, mKast protein expression decreased to approximately 70% in heterozygous mutants compared to that in the WT (Fig. 1E). RNA-seq analysis of the MBs and OLs of WT and mutants revealed that heterozygous mutants exhibited gene expression profiles more similar to those of homozygous mutants than those of the WT (Fig. S2). These results suggest that *mKast* function depends on its product quantity, which could explain the partially similar defects observed in heterozygous and homozygous mutants.

Nevertheless, *mKast* homozygous mutant workers showed a lower survival rate when reared in an indoor colony than heterozygous mutants (Fig. 6D). Worker honey bees forage efficiently by learning multimodal floral stimuli (*38*), and measure flight distances by using an optical flow during foraging flight (*58*). Considering that mKCs are active during foraging flight (*19*), *mKast* likely plays an important role in the brain function to return their hives and thus its knockout mutants result in a lower survival rate when reared in an indoor colony. However, since motion sensing is a general function of insect brains (*59*), and the learning and memory ability of multiple sensory information has been reported even in *Drosophila* (*60*, *61*), these brain functions are not specific to honey bees or other aculeate species. It might be possible that neural connections of mKCs and/or certain OL neurons in honey bees are different from those in other insects such as *Drosophila*, or that the acquisition of selective expression of *mKast* in the adult brain has modified neural functions in these neurons in honey bees.

The present study suggests that *mKast* is involved in the associative learning of bimodal sensory information, but not that of either single olfactory or visual information (Fig. 3 and Fig. S7). In contrast, *CaMKII* is selectively expressed in lKCs in MBs (*62*) and functions in long-term memory in the olfactory PER conditioning paradigm (*18*). Therefore, lKCs and mKCs may be involved in the learning and memory of distinct sensory information and/or learning and memory based on distinct intracellular signaling. mKast IHC signals were detected in the ventral region of the vertical lobe layer corresponding to the basal rings (Fig. 2), suggesting that KCs other than mKCs extend their dendrites into the basal ring and project to the most dorsal layer in the vertical lobe. Based on previous Golgi impregnation (*34*) and IHC of phosphorylated cAMP-response element-binding protein (*63*), this is most likely sKCs. Considering that both sKCs and a part of mKCs are active during foraging flight, and that sKCs are possibly related to lipid metabolism in foraging workers (*22*), mKCs may be involved in the learning and memory of multiple sensory information, while sKCs may be involved in the regulation of the physiological state in foragers. During the evolution of three KC subtypes in aculeate species through functional segregation and divergence from a single KC type in ancestral hymenopteran species (*64*), the acquisition of these distinct mKC functions might have contributed to the evolution of homing ability or high cognitive abilities in aculeate hymenopteran species.

## Materials and Methods

### Animals

European honey bee (*Apis mellifera*) colonies were purchased from Rengedo Beekeepers (Saga, Japan) and Kumagaya Beekeeping Company (Saitama, Japan) and maintained at the University of Tokyo (Tokyo, Japan). Colonies containing genome-edited bees were maintained inside a laboratory room (called the “flight room”) as describe previously (*65*). This was set up per the experimental design using genetically modified organisms approved by the University of Tokyo. The number of workers declined rapidly under indoor rearing conditions and the condition of the indoor colonies tended to deteriorate. Additionally, in some colonies, mosaic queens were made to lay only unfertilized eggs to produce drones. Therefore, comb frames that contained many worker larvae and pupae collected from colonies normally reared outside, or newly emerged adult workers emerged from these frames in an incubator were supplied to colonies set in the flight room whenever necessary to maintain these colonies.

### Production of homozygous mutant workers

Fertilized eggs were collected within 1.5 h after oviposition by confining a queen in a small plastic cage with a honeycomb structure. The eggs were transferred onto a glass slide using a thin paintbrush immersed in ethanol, and injected at their anterior sides with sgRNA targeting *mKast* (200 ng/µL, same sequence as used in the previous study) (*65*) and Cas9 protein (400 ng/µL, PNA Bio) using a glass capillary (Drummond) with its tip broken into a diameter of approx. 5 µm and a microinjector FemtoJet4i (Eppendorf) under an inverted microscope TS100 (Nikon) with the same injection conditions as previous studies (*65*, *66*). The injected eggs were incubated in a humid chamber containing saturated CuSO_4_ at 34 °C. The production of mosaic queens and mutant drones, and collection of semen from drones were conducted as previously described (*65*, *66*). After each queen emerged from the queen cell, one entire forewing was cut with fine scissors and used for genotyping as described below. Mosaic queens that had a high mutation rate (see the genotyping section for details) were used hereafter. After genotyping each drone from which semen was collected, mosaic queens, which were produced in another injection experiment, were artificially inseminated with semen derived from mutant drones using equipment for artificial insemination of honey bees (Schley) (*65*, *67*). They laid fertilized eggs that grew into mutant workers in colonies set in the flight room. Semen from mutant drones was mixed with semen from WT drones, when necessary, and used for the artificial insemination of mosaic queens to let these queens lay fertilized eggs that grew into both heterozygous and homozygous mutant workers.

### Genotyping

Genomic DNA extraction from tissues (one forewing of queens, hind legs of drones, and antennae or hind legs of workers) stored at −20 ℃ after collection, polymerase chain reaction (PCR) to amplify the genomic region around the sgRNA target site, and the sequencing the PCR products were performed as described previously (*65*). Sequence data were processed using Inference of CRISPR Edits (ICE) analysis (*68*) to calculate knockout (KO) scores for mosaic queens or to identify nucleotide insertions and deletions (indels) of both alleles of mutant workers around the sgRNA target site.

The off-target candidate sequences in the honey bee genome were selected using Cas-OFFiner (bulge size ≤ three nucleotides) (*30*). The sequences of the selected off-target candidates were determined by sequencing the PCR products amplified using primers designed near the selected off-target candidates (Table S1) and genomic DNA extracted from WT and mutant workers.

### Olfactory conditioning of PER

Olfactory conditioning of the PER was performed as described previously (*69*), using WT or mutant workers over two weeks old which were outside the colony were collected in a flight room. For the memory test, individual bees were placed in a draft chamber, familiarized for 10 s, and presented with CS or 2-octanol as a novel odor (NOd) for 5 s. After a 10-min interval, another odor, that is, CS or NOd, different from the first, was presented in the same manner. The proportion of bees exhibiting PER when only CS was presented during training or when odors were presented in the memory tests was calculated for each genotype.

### Visual conditioning of PER

Workers over one week old were collected from outside a colony set in a flight room, anesthetized, and fixed on plastic tips with masking tape. Following a previous protocol for visual conditioning of the PER with harnessed honey bees (*70*), both antennae were cut from the base of the scapus using fine scissors. This allowed the harnessed bees to robustly learn visual stimuli. After being fed with 30% (w/w) sucrose until satiation, they were placed in the dark at 25 °C for 20–24 h. Bees were placed in front of a monitor and familiarized for 5 min, and a colored square (7 × 7 cm, blue or green) with a black background was presented on the LED monitor (BenQ, GW2283; 60 Hz refresh rate) for 7 s as a CS. Four seconds after CS presentation, 50% (w/w) sucrose solution (US) was touched on the tip of the proboscis to induce PER and fed for 3 s. This trial was repeated 10 times at 2-min intervals. After performing the memory test 1 h after the end of the training, the bees were fed with 30% (w/w) sucrose solution until saturation and then kept in the dark at 25 °C until the memory test was conducted 24 h after the end of the training. For the memory test, the bees were placed in front of the monitor and familiarized for 5 min, and the CS or a novel color (NCo, blue or green, different from the color used for training) was presented for 7 s. After a 2-min interval, another color that was different from the first was presented in the same manner. The proportion of bees exhibiting PER when only CS was presented during training or when colors were presented in the memory tests was calculated for each genotype.

### Bimodal conditioning

The detailed structure of the apparatus created for this bimodal conditioning was explained in Fig. S5. The combined set of the arena and the transparent slide with eight Petri dishes containing odors (linalool or 2-octanol) and US (+: 50% sucrose, −: 1M NaCl) was placed on a monitor that presented blue or green colors to divide the arena into four quadrants, in which each color was alternately displayed on the monitor in each quadrant (Fig. S5B). Relative light intensities of blue and green displayed on the monitor were adjusted to ensure that bees stayed in each color area for the same amount of time when only color was presented (Fig. S5E). A bee inside the arena could perceive the colors on the monitor through the transparent slide, and the odors or combinations of odors and USs through the holes at the bottom of the arena. By rotating the slide and changing the color on the monitor, the positions of the combinations of colors and odors (CSs) and those of US+/− in the four areas of the arena could be changed.

The training was conducted as follows (Fig. S5F). Workers over one week old were collected outside the colonies set in the flight room and confined in 5 mL tubes. Then, they were fed with 8 µL of 30% sucrose and incubated in the dark at 25 °C for approximately 30 min. Each bee was placed in the arena and familiarized with the monitor displayed in red for 3 min. During familiarization, the bee could not perceive either the CS or the US because the position of the four holes at the bottom of the arena did not match the position of the holes on the slide. By rotating the slide clockwise by 22.5° at the same time as the colors (blue or green) were displayed on the monitor, each of the four holes in the arena presented odors, and each of the four areas of the arena presented a distinct CS (green and linalool / blue and, linalool / green and 2-octanol / blue and 2-octanol) to the bee in the arena. After the CSs were presented for 5 s, the slide was rotated 45 ° clockwise. At this time, each area of the arena presented a combination of US+ (green and linalool) or US− (other CSs) and the CS until the bee licked the US+ for 3 s. The slide was then rotated clockwise by 22.5°, and the color turned red. Bees that did not respond to US+ or required more than 30 min to reach US+ were excluded from subsequent analyses. This training trial was repeated eight times, rotating the positions of the CSs and USs in the arena clockwise by 90° in each trial. To establish the experimental paradigm, an unpaired group was prepared in which the CSs and USs were presented separately. For a trial of the unpaired group, after CSs were presented for 5 s, the slide was rotated clockwise by 22.5°, and the color turned red. After 30 s, the slide was rotated by 22.5° to present USs without CSs (in red light) until the bee licked US+ for 3 s. Then, the slide was rotated 22.5° clockwise and proceeded to the next trial after a 30 s interval. After the training, bees were confined in 5 mL tubes, fed with 8 µL of 30% sucrose, and kept in the dark at 25 °C. Bees were fed with 20 µL of 30% sucrose solution before being kept overnight, and with 8 µL of 30% sucrose solution approximately 20 h and 23 h after the end of the training, respectively. The memory test was performed approximately 24 h after training (Fig. 3B). For the memory test, each bee was placed in the arena and familiarized for 3 min without the CSs and USs (in red light). Then, by rotating the slide clockwise by 22.5°at the same time as the monitor displayed green or blue colors, CSs were presented to the bee for 15 s. The movement of the bees in the arena was video-recorded at five frames per second (fps) with a resolution of 1024 × 768 using a Raspberry Pi camera module (Kumantech) above the arena and tracked using DeepLabCut. A preference index for the CS area conditioned with US+ (a combination of linalool and green) was calculated based on the time spent in each area of the arena as follows: time spent in the CS area minus time spent in other areas divided by the total time.

### Recording antennal responses to motion stimuli

Workers over one week old were collected and anesthetized on ice. Subsequent experiments were performed as previously described (*49*). Briefly, a bee was set between two monitors, familiarized for at least 3 min, and then alternately presented with each of the vertical or horizontal motion stimuli twice with an interstimulus interval of approx. 10 s (antennal movements during this period were ‘spontaneous movements’). The antennal movements of the bee were recorded by camera set above the bee during the experiments.

### Automatic tracking using DeepLabCut

The position of bees in the arena in the bimodal conditioning assay and the antennal movements in the experiments for antennal responses to motion stimuli were tracked using DeepLabCut software (DeepLabCut GUI v2.2.2) (*71*). The antennal movements were tracked as previously described (*49*). For bimodal conditioning, 200 frames were extracted from videos of 20 individuals (10 frames per individual) to create a training dataset for each experiment. The following points were manually labeled on each extracted frame: the center of the arena, the boundaries of the four areas, the Petri dish containing US+, and the anterior and posterior edges of the bee. The network was trained using these data with default parameter settings (ResNet50, 500,000 iterations) and was used to track the position of bees and four areas in the arena in each video.

### Measurement of survival rate

Comb frames containing WT and mutant pupae were set in an incubator at 34 °C, and workers that emerged within one day were collected. For rearing in a plastic cage, mutant (approximately 20 individuals) and WT (approximately 10 individuals) workers within 1 d old were introduced into the plastic cage (95 × 55 × 110 mm) with a piece of honeycomb (6 cm × 7.5 cm) inside and kept in the dark at 25 °C, mimicking the dark conditions inside a hive. Approximately 30 workers of unknown age collected from a normal colony were also introduced into the cage and allowed to care for the newly emerged workers. Bees were fed *ad libitum* with honey and water from 5 mL tubes with small holes set at the bottom and with pollen substitute (Kumagaya Beekeeping Company, Japan) from a 1.5 mL tube inserted near the bottom of the cage. Before caging, one wing of each mutant and WT worker was collected for genotyping, and workers were marked on their dorsal thoraxes with two different colors using a Posca aqueous pen (Mitsubishi, Japan) to identify each individual. For rearing in a flight room colony, workers emerged from same comb frame within one day were separated into two groups: some newly emerged mutant workers were subjected to genotyping, while others were marked on their thorax differently and introduced into the colony in the flight room. After more than two weeks, live marked individuals were collected inside and outside the hive and genotyped. Individuals who died on the floor were removed.

### RNA-seq analysis

Total RNA was extracted from the MBs or OLs of WT and hetero- and homozygous mutant workers one week after emergence using a Direct-zol RNA kit (ZYMO Research). After library preparation with poly-A selection using the MGIEasy Fast RNA Library Prep Set (MGI Tech), RNA-seq was performed using a DNBSEQ G-400 (MGI Tech). The adapter sequences in the RNA-seq reads were trimmed using Trimmomatic (*72*), and the reads were mapped onto the honey bee genome Amel_HAv3.1 using HISAT2 (*73*). Transcripts count matrixes of genes were obtained using StringTie (*74*), and differentially expressed genes (DEGs) were identified using the R package TCC (version 1.32.0) (*75*). Enrichment analysis was conducted using Metascape (*76*) with DEGs in MB and OL, which were converted to *Drosophila melanogaster* orthologs in advance.

### Immunoblotting

After workers were anesthetized on ice, their brains were dissected with fine scissors and homogenized in 100 μL of buffered insect saline (20 mM Tris-HCl, pH 7.4, containing 130 mM NaCl, 5 mM KCl, 1 mM CaCl_2_, and protease inhibitor cocktail [Roche]). They were centrifuged at 10,000× *g* for 10 min, and the supernatant was collected. Four microliters of the protein solution were denatured in sample buffer (50 mM Tris-HCl, pH 6.8, containing 10% glycerol, 2% SDS, 0.002% bromophenol blue, and 3% 2-mercaptethanol), and electrophoresed in an SDS-polyacrylamide gel (5% for stacking, 10% for separation). Immunoblot analysis was performed as described previously (*65*).

### Immunohistochemistry

After workers were anesthetized on ice, their brains were dissected in phosphate-buffered saline (PBS; 137 mM NaCl, 2.7 mM KCl, 8.1 mM Na_2_HPO_4_, 1.5 mM KH_2_PO_4_), and fixed in PBS containing 4% paraformaldehyde for 2–5 d at 4 °C with gentle rotation. The brains were transferred into PBS containing 30% sucrose and rotated for 2–3 h at room temperature, then embedded in 5% agarose in PBS containing 20% sucrose, frozen in liquid nitrogen, and stored at −80 ℃. The brains were sliced into 25 µm-thick sections using a cryostat (Leica) and air-dried at room temperature for at least 2 h. After treatment with PBS for 10 min, 2% TrironX-100 in PBS (PBST) for 10 min, and twice with 0.2% PBST for 10 min twice, heat-induced antigen activation was performed as follows: slides were immersed in a 500 mL beaker containing 400 mL of 10 mM citrate buffer (1.15mM citric acid, 8.85mM sodium citrate, pH 6.0) at 95 °C for 10 min, then the beaker was transferred to room temperature (*77*). After 10 min, the beaker was immersed in water at room temperature for 20 min. After washing with 0.2% PBST, slides were treated with 0.2% PBST containing 2% normal goat serum for 1 h at room temperature. The slides were treated with primary antibody solution (rabbit anti-FMRFamide polyclonal antibody [1:1000, Enzo Life Sciences, BML-FA1155], mouse anti-SYNORF1 monoclonal antibody [1:20, Developmental Studies Hybridoma Bank, 3C11] (*78*), guinea pig anti-mKast polyclonal antibody [0.5 µg/mL, produced in the previous study] (*26*)) overnight at 4 °C with shaking. After washing with PBS five times, the slides were treated with a secondary antibody solution (goat anti-mouse IgG antibody conjugated with Alexa Fluor 488 [1:400, Invitrogen, A11001], goat anti-rabbit IgG antibody conjugated with Alexa Fluor 546 [1:400, Invitrogen, A11071], and goat anti-guinea pig IgG antibody conjugated with Alexa Fluor 546 [1:400, Invitrogen, A11074]) for 2 h at room temperature in the dark. Then, the slides were treated with 10 µg/mL Hoechst33342 (Lonza) in PBS for 15 min at room temperature, followed by washing with PBS five times, replacement with 60% glycerol for 30 min, and mounting in 80% glycerol. Fluorescence images were captured using a fluorescence microscope BZ-X800 (Keyence).

### *In situ* hybridization

Workers were collected near the entrance of outside colonies, and after anesthetization on ice, their brains were dissected in insect saline, embedded in O.C.T.compound (Sakura Fintek, Japan), frozen on dry ice, and stored at −80 °C. *In situ* hybridization (ISH) was performed using 10 µm-thick sections sliced in a cryostat as described previously (*26*). The cDNA fragment of *Sox102F* was amplified with primers (Fw: 5′-CACCACACGAACACATCCTC-3′, Rv: 5′-GCTATCCGAGATGATCCTGC-3′) and One*Taq* DNA polymerase (NEB)using cDNA reverse-transcribed with SuperScrptIII (Invitrogen) as a template and cloned into pGEM-t easy vector (Promega). The cDNA fragment of *mKast* was cloned into pGEM-t as described in a previous study. RNA probes were synthesized using these cloned vectors, as described previously (*26*) and stored at −80 °C until use.

### Comparative analysis of honey bee and *Drosophila* OL scRNA-seq data by MetaNeighbor

The scRNA-seq data of the OLs of honey bee foragers (DRA014752) (*22*) and adult *Drosophila* (GSE142787) (*44*) from previous studies were used in this study. The honey bee OL scRNA-seq data were reanalyzed using the R package Seurat (version 4.2.0) (*79*). The data were preprocessed as described in a previous study. However, cells with nFeature_RNA < 6000 were used in this study. After normalization with the LogNormalize option, cells were clustered with the parameters npcs = 50 and resolution = 1.5. The clusters and annotations of adult *Drosophila* OL scRNA-seq data were the same as those used in the previous study (*44*). A pretrained MetaNeighbor model was created from the *Drosophila* data using MetaNeighbor (*45*). Using genes with reciprocal best hits between the honey bee and *Drosophila*, the area under the receiver operating characteristic curve (AUROC) scores between clusters of the honey bee and *Drosophila* were calculated using the pre-trained model. Cytoscape (*80*) was used to plot the correspondence between the clusters with high AUROC scores.

### Statistical Analysis

All statistical analyses were performed using the R software environment. A detailed description of the statistics, including individual data points, and p-values are provided in the figure legends. All tests were 2-tailed.

### Data and materials availability

The raw data obtained from the RNA-seq analyses were deposited in the DNA Data Bank of Japan Sequence Read Archive under accession number PRJDB19785.

## Author contributions

Conceptualization: HK, TK

Methodology: HK

Investigation: HK

Visualization: HK

Funding acquisition: HK, TK

Project administration: HK, TK

Supervision: TK

Writing – original draft: HK

Writing – review & editing: HK, TK

## Supplementary Figures

**Fig. S1.**
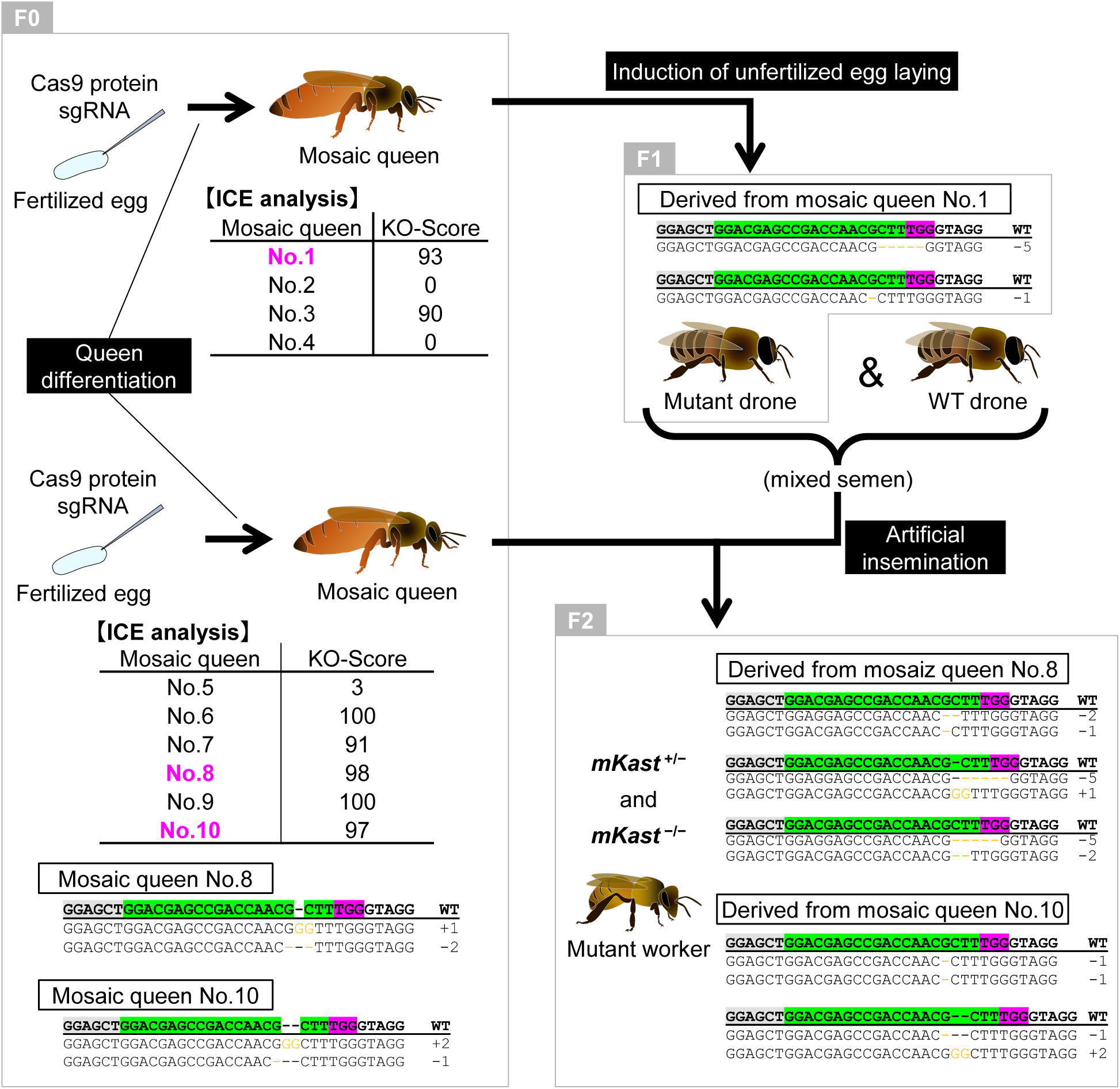
Details of the process to produce *mKast* homozygous mutant workers. The process to produce mutant workers (F2). Results of a representative experiment are shown. Tables in the left panel show the mutation rates (KO scores) in queens estimated by ICE analysis using the genomic DNA extracted from one forewing of each mosaic queen (F0; Nos. 1-10). Sequences show the indels detected at the sgRNA target site of some mosaic queens (F0), mutant drones (F1), and homozygous mutant workers (F2) by genotyping using ICE analysis. Letters marked in green, magenta, and gray in WT sequence indicate the sgRNA target site, PAM sequence, and exon of *mKast*, respectively. Orange letters and dashes indicate the inserted and deleted nucleotide sequences, respectively. Mosaic queens with high KO score (Nos. 1, 8 and 10) were selected and used to produce the next generation. In this experiment, mosaic queens No. 8 and 10 (F0) were used for artificial insemination with the semen of mutant drones (F1) produced by mosaic queen No. 1 (F0). *mKast* homozygous mutant workers (F2) possessed indels corresponding to those in the mosaic queens (F0) and mutant drones (F1) used for artificial insemination. For example, indels detected in homozygous mutant workers (F2) derived from mosaic queen No. 8 (−2 / −1, −5 / +1, and −5 / −2) corresponded to either of them in mosaic queen No. 8 (+1 / −2) and that in mutant drones (−5 or −1).

**Fig. S2.**
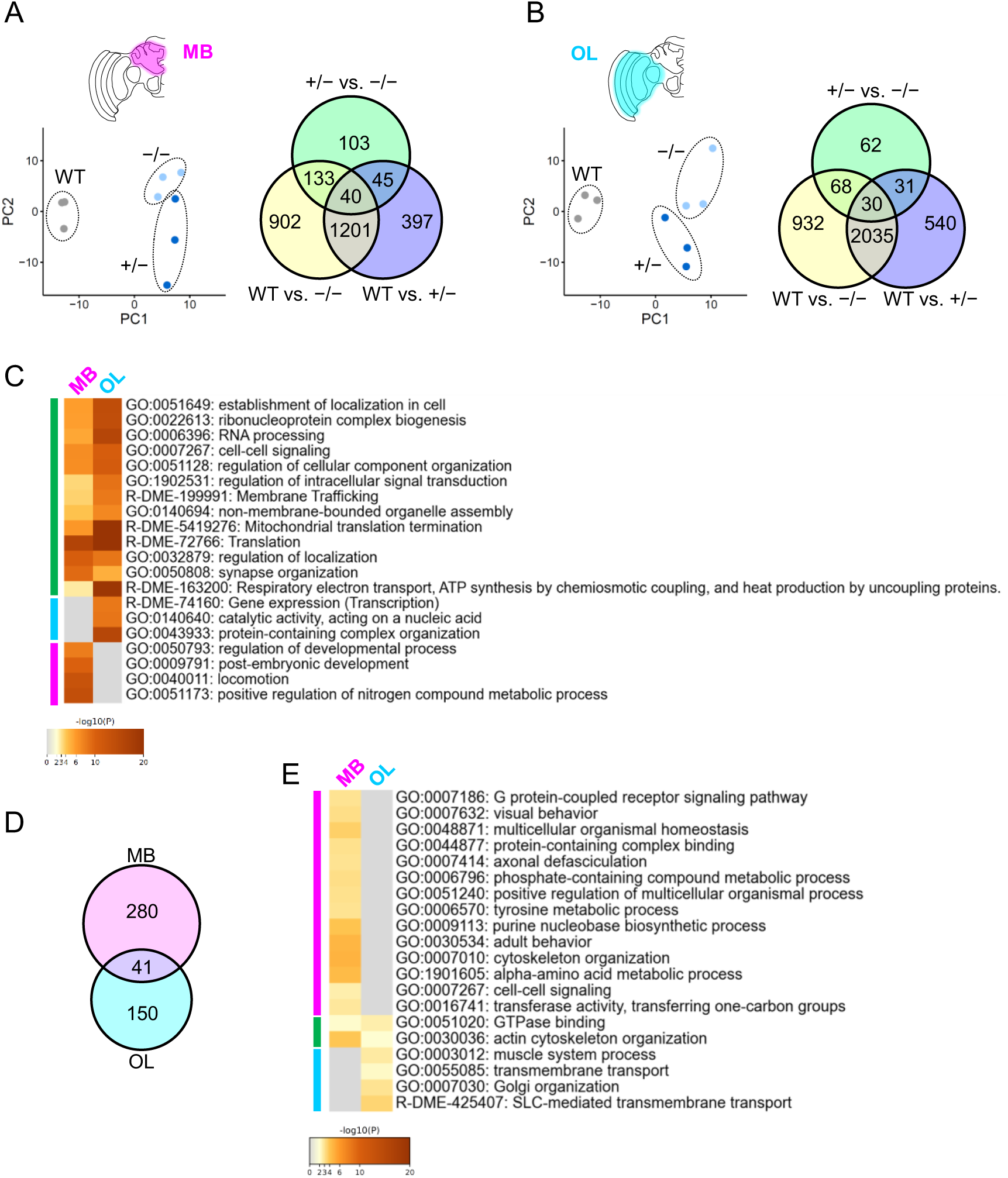
RNA-seq analysis of the MBs and OLs of WT and mutant workers. (A, B) Summary of RNA-seq results for the MBs (A) and OLs (B) of adult workers of each genotype (WT, +/−, −/−). Left graphs show results of principal component analysis of all RNA-seq samples. Right pie charts show the number of DEGs (p < 0.01) between the three genotypes. (C) GO analysis of DEGs (q < 0.05) among genotypes in the MBs and OLs using Metascape. Green, light blue, and magenta bars indicate GO terms detected in the DEGs of both MBs and OLs, OLs, and MBs, respectively. (D) Pie chart showing the number of DEGs (p < 0.01) detected between hetero- and homozygous mutants in the MBs and OLs. (E) Results of GO analysis of DEGs (D) between hetero- and homozygous mutants in the MBs and OLs using Metascape. Magenta, green, and light blue bars indicate GO terms detected in the DEGs of MBs, both MBs and OLs, and OLs, respectively.

**Fig. S3.**
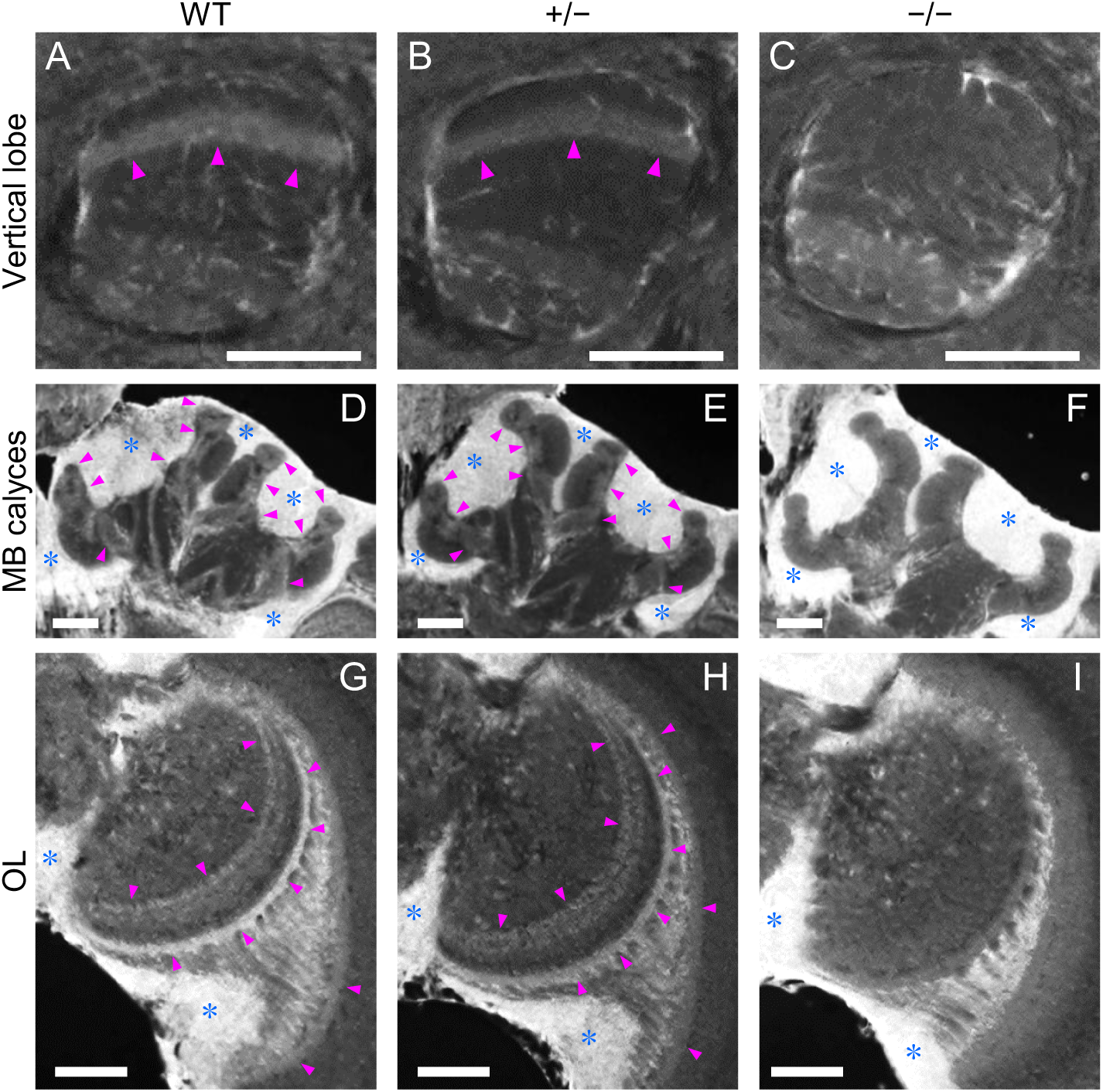
Localization of mKast protein in the brains of WT and mutant workers. mKast IHC results in the vertical lobes (A–C) and calyces (D–F) of the MBs, and the OLs (G–I) in the brains of WT (A, D, G), heterozygous mutants (B, E, H) and homozygous mutants (C, F, I). Magenta arrowheads indicate the mKast IHC signals and blue asterisks indicate non-specific fluorescent signals in the cell body region. Scale bar, 100 µm.

**Fig. S4.**
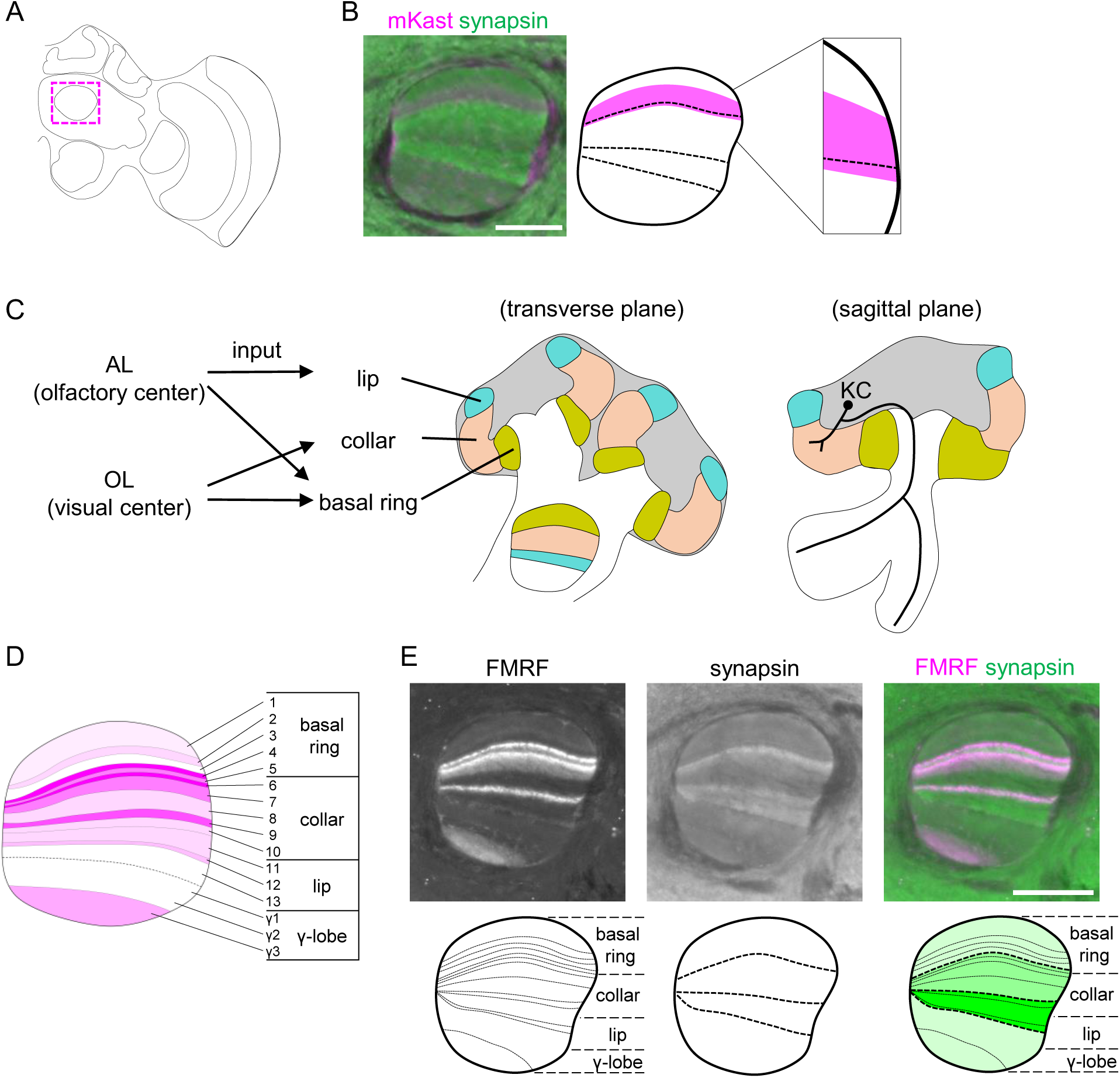
Correspondence between MB calyceal compartments where sensory information inputs and mKast ISH signal in the MB vertical lobe. (A) A schematic diagram of anterior section of an adult brain. The magenta dotted box indicates the location of the vertical lobe shown in panels B–D. (B) Results of IHC using anti-mKast and synapsin antibodies in WT and a schematic diagram of the mKast IHC signal in the vertical lobe (reproduced from Fig. 2B), with an enlarged schematic diagram added. (C) Schematic diagrams of MB in the transverse and sagittal planes (right), and input patterns from primary sensory centers to the MB calyceal compartments (left) reproduced from Strausfeld (2002). (D) A schematic diagram of the vertical lobe reproduced from Strausfeld (2002). The magenta intensity pattern indicates IHC signals using anti-FMRFamide antibodies. The layers of projection areas from the MB calyceal compartments are shown. (E) Results of IHC with anti-synapsin and FMRFamide antibodies in WT. Schematic diagrams below the images indicate the layered signal pattern of each IHC. The schematic diagram below the merged image shows the correspondence between the layered structure by synapsin IHC (indicated by green scales) and projection areas from calyceal compartments drawn based on the signal pattern of FMRFamide IHC (dotted line). Scale bar, 100 µm.

**Fig. S5.**
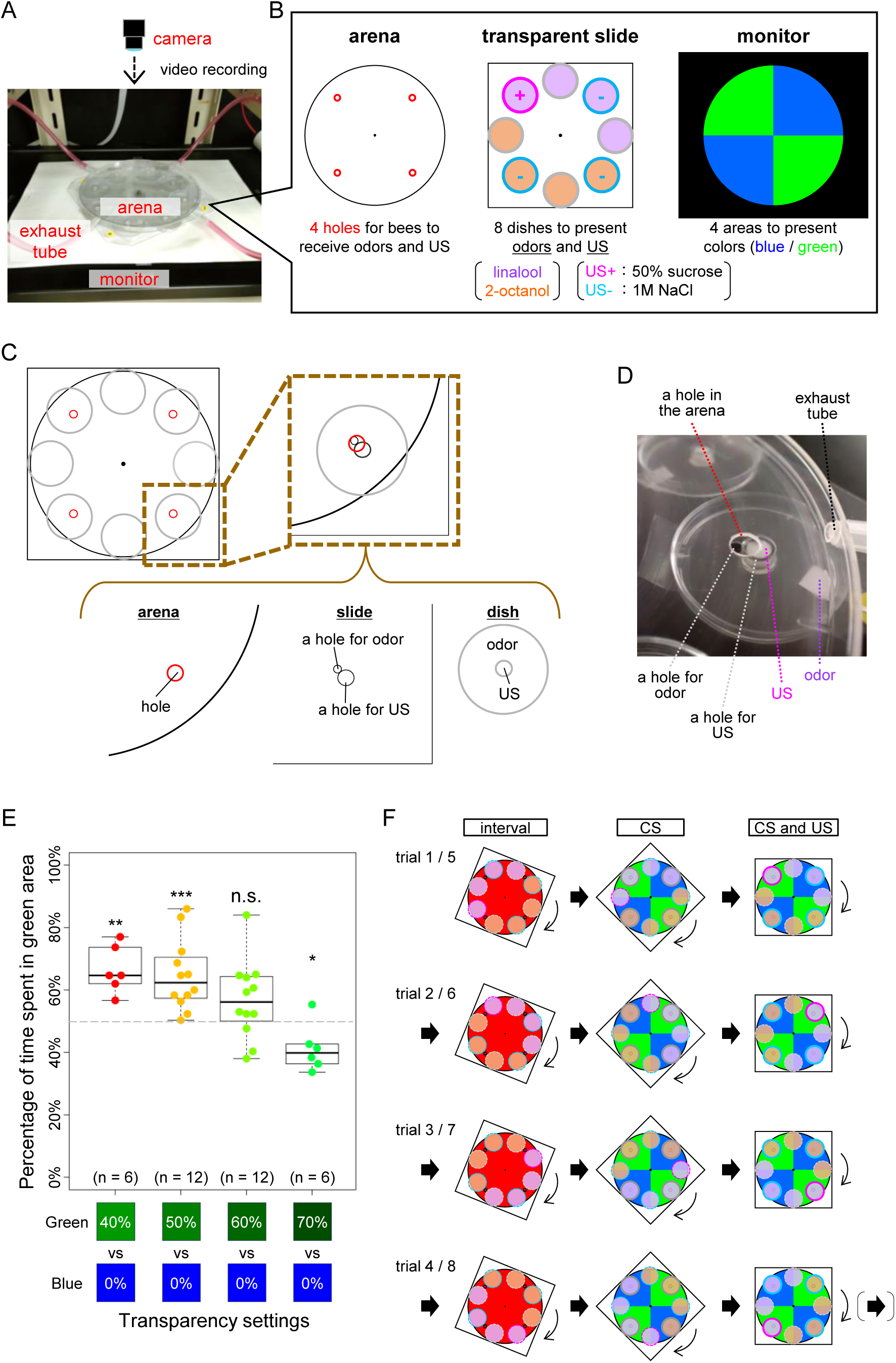
Experimental setup for bimodal conditioning assay. (A) Experimental setup in which bimodal conditioning was performed. A transparent slide (15 × 15 cm) is placed between the monitor and the arena (15 cm in diameter and 7 mm in height), and video is recorded by a camera from above. Exhaust tubes are connected to the arena to ventilate inside. (B) Schematic diagram of each component of the experimental setup: the arena (left), transparent slide (middle), and monitor (right). The arena has four holes (red circles) through which a bee receives odors and US. Eight small transparent Petri dishes (38 mm in diameter and 5 mm in height) are placed in a circular arrangement underneath the transparent slide. The eight Petri dishes on the transparent slide contained a piece of filter paper (5 × 5 mm) impregnated with an odor solution (46.8% linalool in mineral oil [purple] in four consecutive dishes or 50% 2-octanol in mineral oil [orange] in another four dishes). One Petri dish containing linalool was supplied with a piece of sponge soaked in 50% (w/w) sucrose as the US of reward (US+), while three other Petri dishes, one of which was filled with linalool and two with 2-octanol were supplied with pieces of sponge soaked in 1M NaCl as the US of punishment (US−). The monitor displays blue or green colors alternatively in four areas of the arena. (C) Details of the assembly of the arena, slide, and Petri dish is shown using one Petri dish as an example. Eight holes of 6 mm in diameter were drilled on the slide at positions corresponding to the center of the eight Petri dishes. An additional hole of 3 mm in diameter was drilled at the inner position adjacent to the larger hole. The circular arena was placed on the slide, and a shaft was attached to the center of the slide and arena to enable the slide to rotate against the arena. Four holes were drilled at the bottom of the arena to make the bees in the arena accessible to the odors and lick USs from dishes beneath the arena through 3-mm and 8-mm diameter holes on the slide, respectively. (D) An image when presenting an odor and US in the Petri dish under the hole at the bottom of the arena through two holes in the slide. A clear plastic cover was placed on the arena to prevent the bees from escaping. The odor presented from the Petri dishes was constantly exhausted through tubes, which were connected to fans, from four locations on the sides of the arena. (E) Adjustment of the balance of blue and green color intensities displayed on the monitor by changing transparency settings of each color. Percentage of time bees spent in each color area was measured. One-sample t-test was used to test whether the percentage of time spent in the green area was different from 50%. *** p < 0.001, ** p < 0.01, * p < 0.05, n.s., not significant. The combination of 60% green and 0% blue, which had no significant differences, was adopted in the bimodal conditioning assay. (F) The position of a transparent slide with eight transparent Petri dishes and four areas (blue or green) displayed on the monitor in each trial of training process during bimodal conditioning paradigm. The purple and orange dishes contain linalool and 2-octanol, respectively. The dish surrounded with magenta dashed or solid lines contains 50% sucrose as a reward, and the dishes surrounded with light blue dashed or solid lines contain 1M NaCl as punishment. In the interval, when the monitor displays red color, the position of the holes at the bottom of the arena and the position of holes on the slide do not match, and so both odors or USs are not presented to the bee in the arena. For CSs presentation, by rotating the slide 22.5° cloclwise, odors are presented from the Petri dish underneath the slide through the holes at the bottom of the arena, and colors are displayed on the monitor. For the presentation of CSs and USs, the slide is further rotated 45° clockwise so that the center of the Petri dishes containing odors and USs are positioned under the hole at the bottom of the arena. In each trial, the position of CSs and the combinations of CSs and USs in the arena is rotated by 90°. See Materials and Methods for more details.

**Fig. S6.**
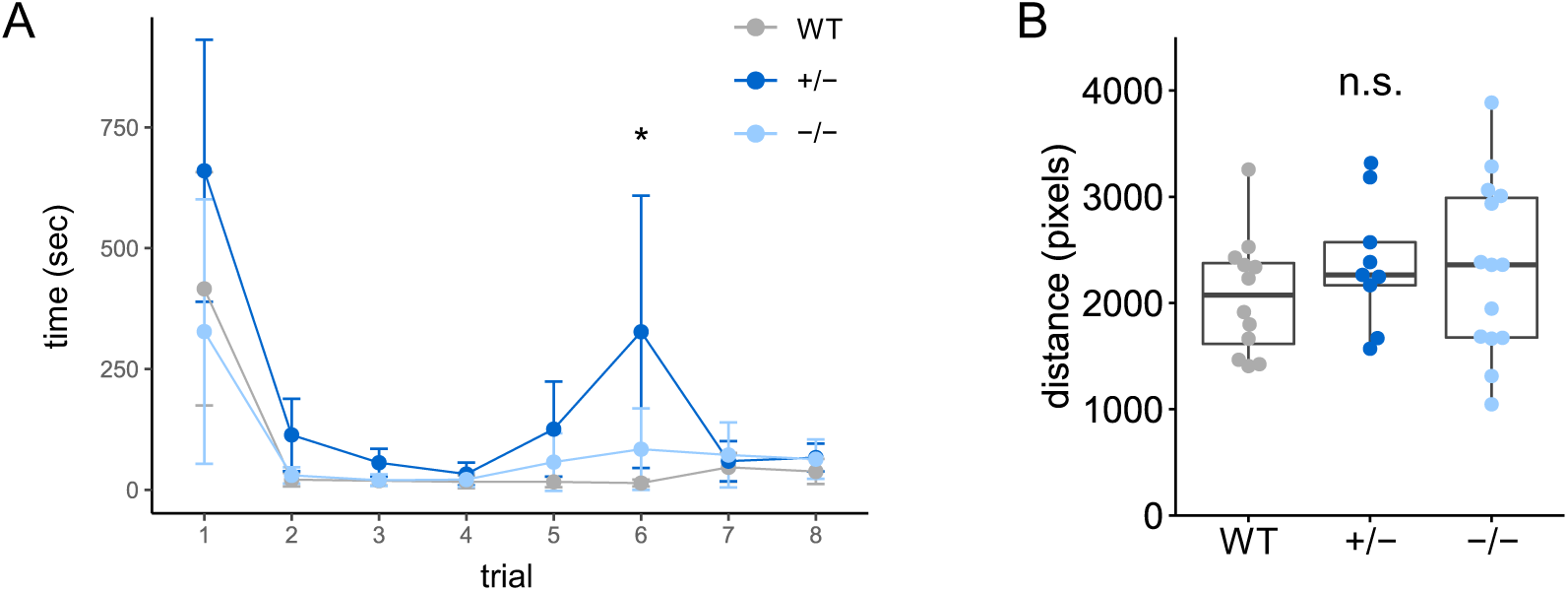
Time to reach US+ in each trial during training and moving distances during memory test of bimodal conditioning assay. (A) Time taken to reach US+ in each trial during training for each genotype. Kruskal-Wallis test, * p < 0.05. (B) Moving distances during memory test (15 s) for each genotype. ANOVA, n.s., not significant. n = 12, 9, and 14 in WT, hetero- and homozygous mutants, respectively.

**Fig. S7.**
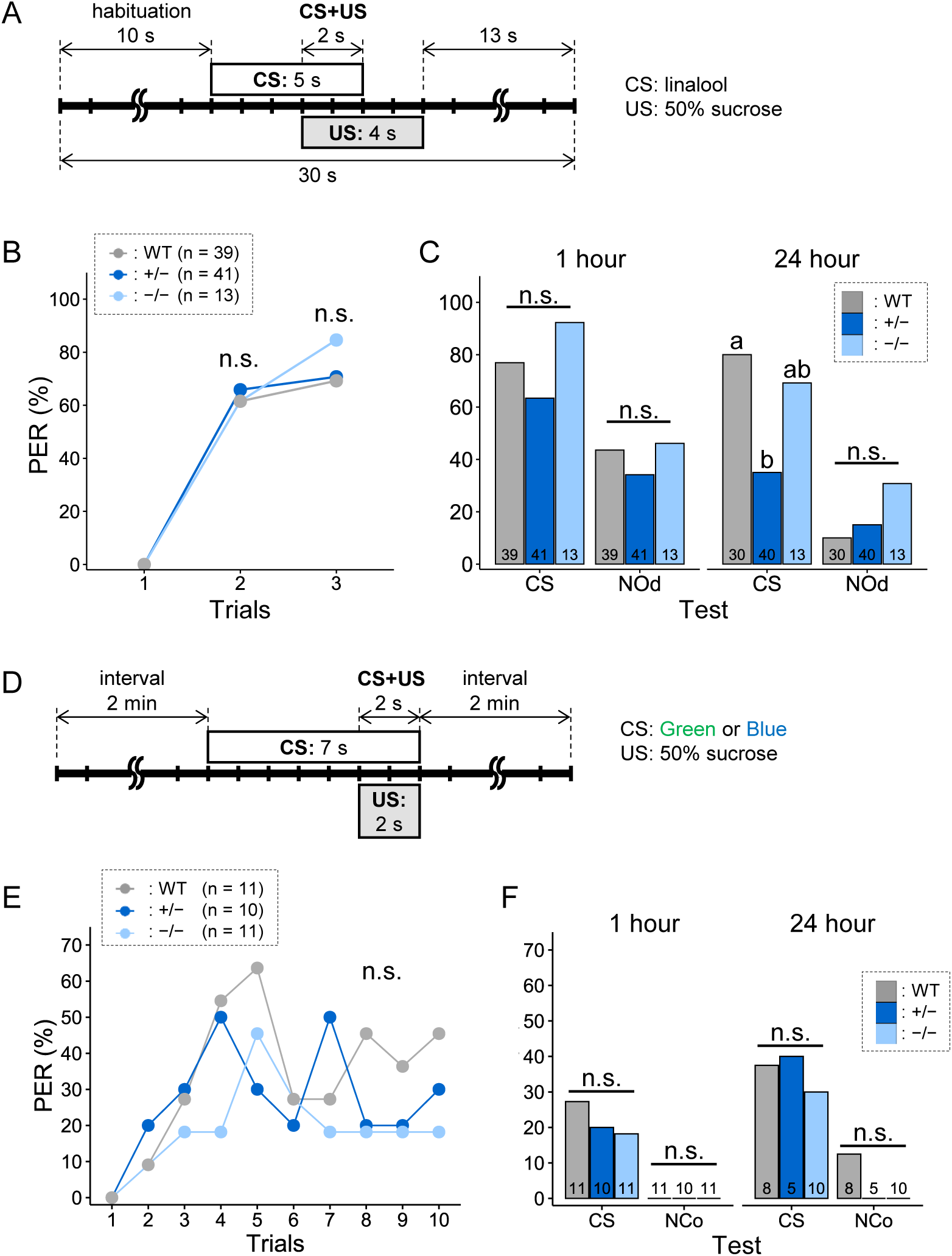
Olfactory or visual PER conditioning in WT and mutant workers. (A) Experimental scheme of each trial in the training of the associative learning of odor (linalool) and PER, in which CS alone was presented followed by US in addition to CS. Trial was repeated three times with 10-minute interval. (B, C) Results of the training (B) and the memory test at 1 h or 24 h after the end of training (C) in the olfactory PER conditioning experiment for each genotype. The percentage of individuals that exhibited PER when only CS was presented are shown. NOd, novel odor (2-octanol). The numbers inside each bar in (C) indicate the number of individuals. Fisher’s exact test with Hochberg correction, different letters indicate significant difference (p < 0.05), n.s., not significant. (D) Experimental scheme of each trial in the training of the associative learning of color (green or blue) and PER, in which CS alone was presented followed by US in addition to CS. Trial was repeated 8 times with 2-min interval. (E, F) Results of the training (E) and the memory test at 1 hr or 24 hr after the end of training (F) in the visual PER conditioning experiment for each genotype. The percentage of individuals that exhibited PER when only CS was presented are shown. NCo, novel color (green or blue, different color from CS). The numbers inside each bar in (F) indicate the number of individuals; Fisher’s exact test with Hochberg correction, n.s., not significant.

**Fig. S8.**
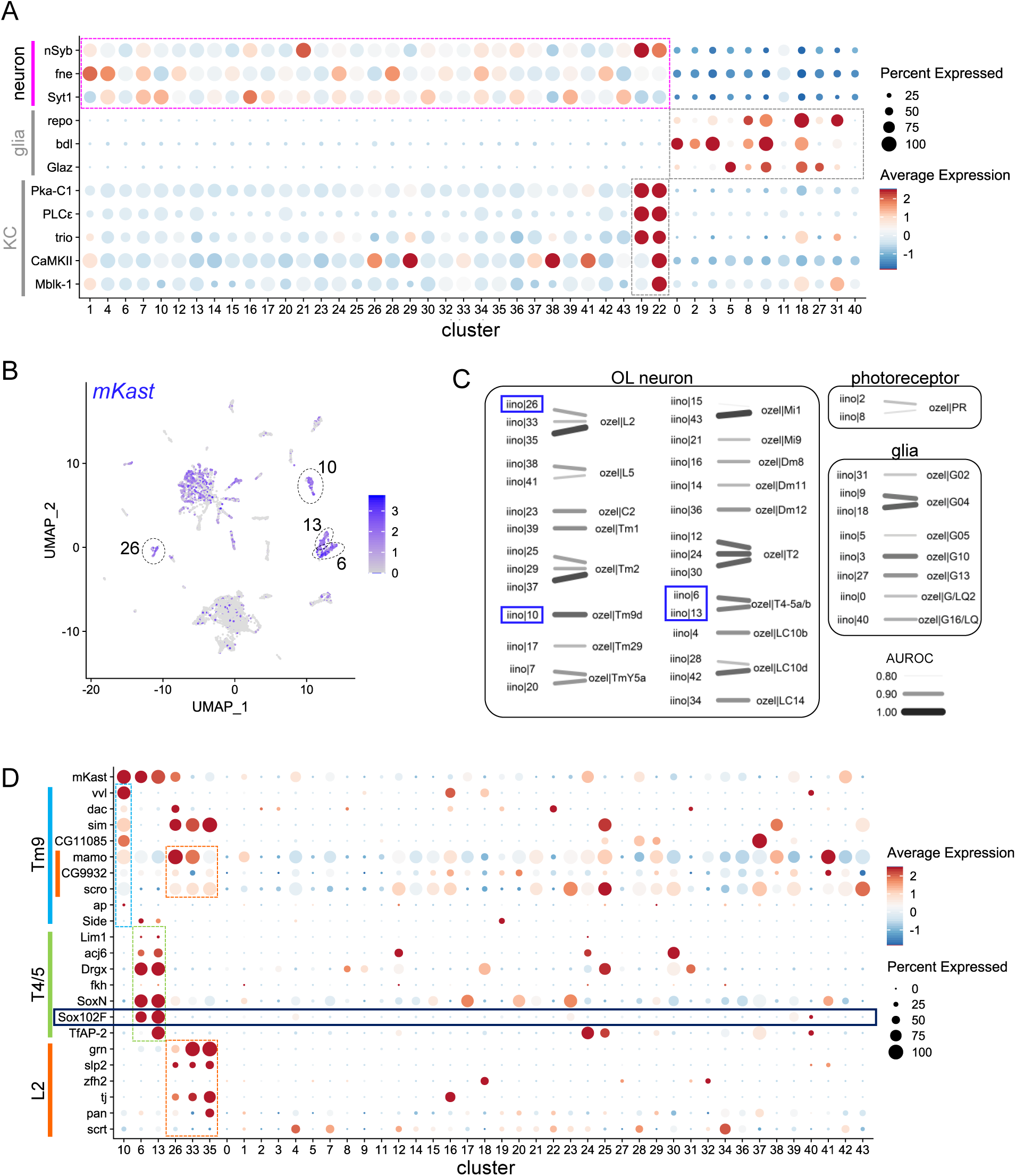
Comparative analysis of OL scRNA-seq data between the honey bee and fruit fly. (A) A dot plot of neural, glial, and KC marker genes in clusters of re-analyzed adult honey bee OL scRNA-seq data (Iino et al., 2023). (B) A feature plot showing *mKast* expression pattern in cells of honey bee OL scRNA-seq data. Clusters with high *mKast* expression level (clusters 6, 10, 13, and 26) are indicated as surrounded by dotted lines. (C) The correspondence of clusters with high similarity (AUROC score > 0.80) between honey bee and *Drosophila* clusters is shown. Only *Drosophila* clusters to which the honey bee OL clusters show the highest AUROC scores are extracted. The density and width of the line connecting the clusters reflect the AUROC score. The violet boxes indicate clusters with high *mKast* expression level. AUROC, area under the receiver operating characteristic curve. (D) A dot plot showing the expression levels of marker genes for *Drosophila* L2, Tm9, and T4/5 neurons in the honey bee OL cell clusters. Light blue, yellow-green, and orange bars indicate marker genes in Tm9, T4/5, and L2 neurons, respectively.

**Fig. S9.**
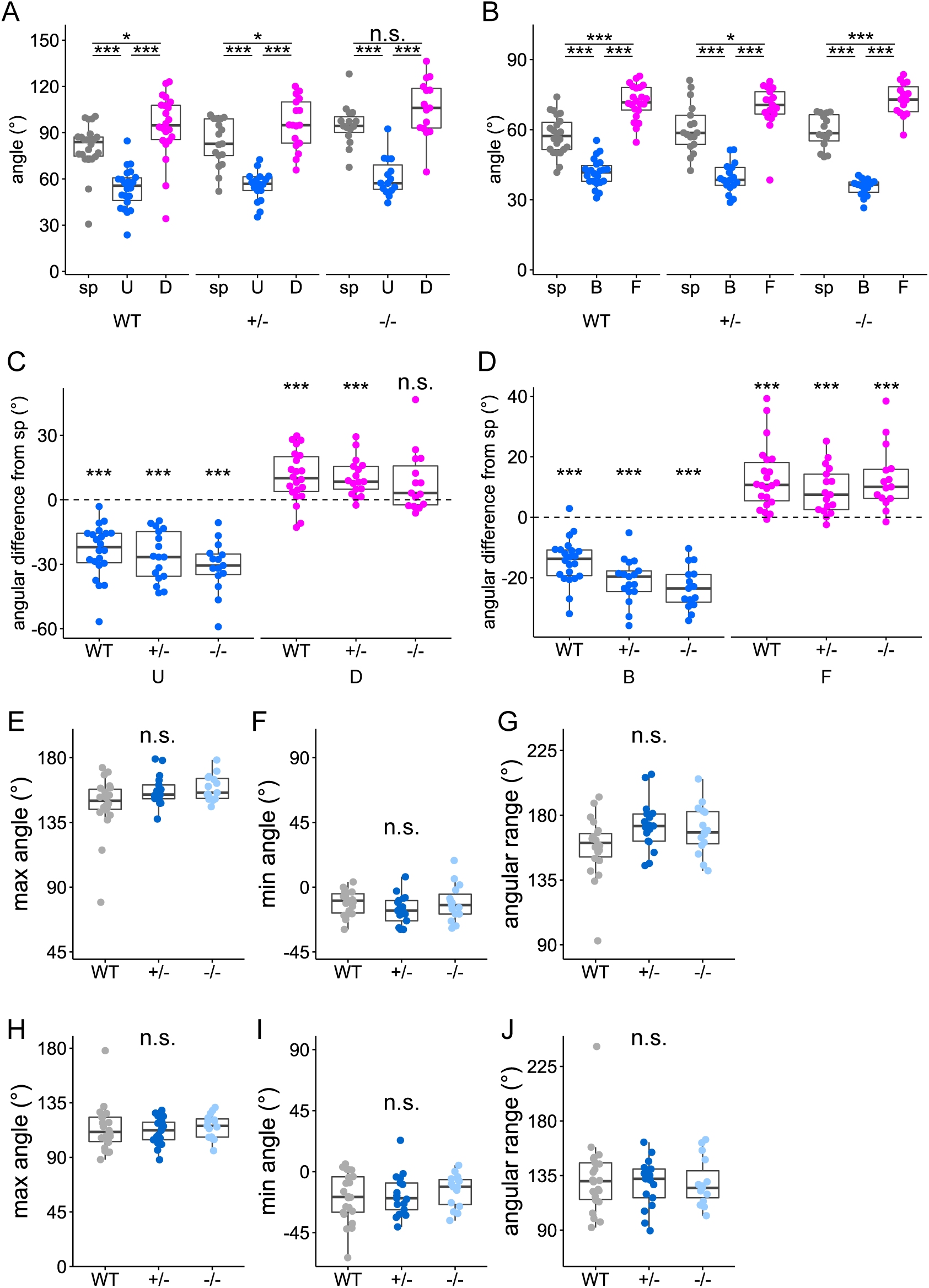
ARs to motion stimuli and angular ranges of antennae in WT and mutant workers. (A, B) Comparison of the angles of the antennae during the presentation of motion stimuli and spontaneous movements in the transverse plane (A) and coronal plane (B) in each genotype. Tukey HSD test, *** p < 0.001, * p < 0.05, n.s., not significant. (C, D) The differences between the angles of the antennae during the presentation of each motion stimulus and during spontaneous movement in the transverse plane (C) and coronal plane (D) are shown for each genotype. One sample Wilcoxon test was used to test whether the angular difference was different from 0°. *** p < 0.001, n.s., not significant. (E–J) Maximum (E, H) and minimum (F, I) angles and angular ranges (difference between maximum and minimum angles) (G, J) of each genotype during spontaneous movement of antennae in the transverse plane (E–G) and coronal plane (H–J). Steel–Dwass test, n.s., not significant. In all graphs in this figure, n = 22, 17, 15 in WT, heterozygous mutants, and homozygous mutants, respectively.

## References

1. J. Korb, J. Heinze, Major Hurdles for the Evolution of Sociality. Annu. Rev. Entomol. 61, 297–316 (2016).

2. T. Clutton-Brock, Social evolution in mammals. Science 373, eabc9699 (2021).

3. M. Anderson, The Evolution of Eusociality. Annu. Rev. Ecol. Evol. Syst. 15, 165–189 (1984).

4. R. S. Peters, L. Krogmann, C. Mayer, A. Donath, S. Gunkel, K. Meusemann, A. Kozlov, L. Podsiadlowski, M. Petersen, R. Lanfear, P. A. Diez, J. Heraty, K. M. Kjer, S. Klopfstein, R. Meier, C. Polidori, T. Schmitt, S. Liu, X. Zhou, T. Wappler, J. Rust, B. Misof, O. Niehuis, Evolutionary History of the Hymenoptera. Curr. Biol. 27, 1013–1018 (2017).

5. M. Collett, L. Chittka, T. S. Collett, Spatial Memory in Insect Navigation. Curr. Biol. 23, R789– R800 (2013).

6. A. Avarguès-Weber, M. Giurfa, Conceptual learning by miniature brains. Proc. R. Soc. B Biol. Sci. 280, 20131907 (2013).

7. E. A. Tibbetts, Visual signals of individual identity in the wasp Polistes fuscatus. Proc. R. Soc. London. Ser. B Biol. Sci. 269, 1423–1428 (2002).

8. M. Giurfa, Honeybees foraging for numbers. *J. Comp. Physiol. A Neuroethol. Sensory, Neural*, Behav. Physiol. 205, 439–450 (2019).

9. K. V. Frisch, A. M. Wenner, D. L. Johnson, Honeybees: Do They Use Direction and Distance Information Provided by Their Dancers? Science 158, 1072–1077 (1967).

10. M. L. Winston, The Biology of the Honey Bee (Harvard University Press, Cambridge, 1987).

11. C. W. Whitfield, A. M. Cziko, G. E. Robinson, Gene expression profiles in the brain predict behavior in individual honey bees. Science 302, 296–299 (2003).

12. G. S. Withers, S. E. Fahrbach, G. E. Robinson, Selective neuroanatomical plasticity and division of labour in the honeybee. Nature 364, 238–240 (1993).

13. T. Zars, M. Fischer, R. Schulz, M. Heisenberg, Localization of a short-term memory in *Drosophila*. Science 288, 672–675 (2000).

14. M. Mizunami, J. M. Weibrecht, N. J. Strausfeld, Mushroom bodies of the cockroach: Their participation in place memory. J. Comp. Neurol. 402, 520–537 (1998).

15. S. M. Farris, S. Schulmeister, Parasitoidism, not sociality, is associated with the evolution of elaborate mushroom bodies in the brains of hymenopteran insects. Proc. R. Soc. B 278, 940–951 (2011).

16. S. Oya, H. Kohno, Y. Kainoh, M. Ono, T. Kubo, Increased complexity of mushroom body Kenyon cell subtypes in the brain is associated with behavioral evolution in hymenopteran insects. Sci. Rep. 7, 13785 (2017).

17. K. Kaneko, S. Suenami, T. Kubo, Gene expression profiles and neural activities of Kenyon cell subtypes in the honeybee brain: identification of novel ‘middle-type’ Kenyon cells. Zool. Lett. 2, 14 (2016).

18. C. Scholl, N. Ku bert, T. S. Muenz, W. Rossler, CaMKII knockdown affects both early and late phases of olfactory long-term memory in the honeybee. J. Exp. Biol. 218, 3788–3796 (2015).

19. K. Kaneko, T. Ikeda, M. Nagai, S. Hori, C. Umatani, H. Tadano, A. Ugajin, T. Nakaoka, R. K. Paul, T. Fujiyuki, K. Shirai, T. Kunieda, H. Takeuchi, T. Kubo, Novel middle-type Kenyon cells in the honeybee brain revealed by area-preferential gene expression analysis. PLoS One 8, e71732 (2013).

20. T. Kiya, T. Kunieda, T. Kubo, Increased Neural Activity of a Mushroom Body Neuron Subtype in the Brains of Forager Honeybees. PLoS One 2, e371 (2007).

21. S. Iino, Y. Shiota, M. Nishimura, S. Asada, M. Ono, T. Kubo, Neural activity mapping of bumble bee (*Bombus ignitus*) brains during foraging flight using immediate early genes. Sci. Rep. 10, 7887 (2020).

22. S. Iino, S. Oya, T. Kakutani, H. Kohno, T. Kubo, Identification of ecdysone receptor target genes in the worker honey bee brains during foraging behavior. Sci. Rep. 13, 10491 (2023).

23. A. Roth, C. Vleurinck, O. Netschitailo, V. Bauer, M. Otte, O. Kaftanoglu, R. E. Page, M. Beye, A genetic switch for worker nutritionmediated traits in honeybees. PLoS Biol. 17, e3000171 (2019).

24. L. Puca, C. Brou, α-Arrestins – new players in Notch and GPCR signaling pathways in mammals. J. Cell Sci. 127, 1359–1367 (2014).

25. C. H. Lin, J. A. MacGurn, T. Chu, C. J. Stefan, S. D. Emr, Arrestin-Related Ubiquitin-Ligase Adaptors Regulate Endocytosis and Protein Turnover at the Cell Surface. Cell 135, 714–725 (2008).

26. A. Yamane, H. Kohno, T. Ikeda, K. Kaneko, A. Ugajin, T. Fujita, T. Kunieda, T. Kubo, Gene expression and immunohistochemical analyses of mKast suggest its late pupal and adult-specific functions in the honeybee brain. PLoS One 12, e0176809 (2017).

27. Z. Chen, I. M. Traniello, S. Rana, A. C. Cash-Ahmed, A. L. Sankey, C. Yang, G. E. Robinson, Neurodevelopmental and transcriptomic effects of CRISPR/Cas9-induced somatic *orco* mutation in honey bees. J. Neurogenet. 35, 320–332 (2021).

28. D. A. De Souza, O. Kaftanoglu, D. De Jong, R. E. Page, G. V. Amdam, Y. Wang, Differences in the morphology, physiology and gene expression of honey bee queens and workers reared in vitro versus in situ. Biol. Open 7, bio036616 (2018).

29. A. N. Mortensen, J. D. Ellis, The effects of artificial rearing environment on the behavior of adult honey bees, *Apis mellifera* L. Behav. Ecol. Sociobiol. 72, 92 (2018).

30. S. Bae, J. Park, J. S. Kim, Cas-OFFinder: a fast and versatile algorithm that searches for potential off-target sites of Cas9 RNA-guided endonucleases. Bioinformatics 30, 1473–1475 (2014).

31. J. E. Garneau, M. È. Dupuis, M. Villion, D. A. Romero, R. Barrangou, P. Boyaval, C. Fremaux, P. Horvath, A. H. Magadán, S. Moineau, The CRISPR/Cas bacterial immune system cleaves bacteriophage and plasmid DNA. Nature 468, 67–71 (2010).

32. K. Zbieralski, D. Wawrzycka, α-Arrestins and Their Functions: From Yeast to Human Health. Int. J. Mol. Sci. 23, 4988 (2022).

33. K. T. Lee, I. K. A. Pranoto, S. Y. Kim, H. J. Choi, N. B. To, H. Chae, J. Y. Lee, J. E. Kim, Y. V. Kwon, J. W. Nam, Comparative interactome analysis of α-arrestin families in human and *Drosophila*. Elife 12, RP88328 (2024).

34. N. J. Strausfeld, Organization of the honey bee mushroom body: Representation of the calyx within the vertical and gamma lobes. J. Comp. Neurol. 450, 4–33 (2002).

35. C. G. Galizia, W. Rössler, Parallel olfactory systems in insects: Anatomy and function. Annu. Rev. Entomol. 55, 399–420 (2010).

36. M. V. Srinivasan, Honey Bees as a Model for Vision, Perception, and Cognition. Annu. Rev. Entomol. 55, 267–284 (2010).

37. J. Erber, T. Masuhr, R. Menzel, Localization of short-term memory in the brain of the bee, *Apis mellifera*. Physiol. Entomol. 5, 343–358 (1980).

38. A. S. Leonard, P. Masek, Multisensory integration of colors and scents: Insights from bees and flowers. *J. Comp. Physiol. A Neuroethol. Sensory, Neural*, Behav. Physiol. 200, 463–474 (2014).

39. D. Thiagarajan, S. Sachse, Multimodal Information Processing and Associative Learning in the Insect Brain. Insects 13, 332 (2022).

40. M. Giurfa, J.-C. Sandoz, Invertebrate learning and memory: Fifty years of olfactory conditioning of the proboscis extension response in honeybees. Learn. Mem. 19, 54–66 (2012).

41. M. Kuwabara, Bildung des bedingten Reflexes von Pavlovs Typus bei der Honigbiene, Apis mellifica. J Fac Hokkaido Uni Ser VI Zool 13, 458–464 (1957).

42. W. Zhang, L. Wang, Y. Zhao, Y. Wang, C. Chen, Y. Hu, Y. Zhu, H. Sun, Y. Cheng, Q. Sun, J. Zhang, D. Chen, Single-cell transcriptomic analysis of honeybee brains identifies vitellogenin as caste differentiation-related factor. iScience 25, 104643 (2022).

43. Q. Li, M. Wang, P. Zhang, Y. Liu, Q. Guo, Y. Zhu, T. Wen, X. Dai, X. Zhang, M. Nagel, B. H. Dethlefsen, N. Xie, J. Zhao, W. Jiang, L. Han, L. Wu, W. Zhong, Z. Wang, X. Wei, W. Dai, L. Liu, X. Xu, H. Lu, H. Yang, J. Wang, J. J. Boomsma, C. Liu, G. Zhang, W. Liu, A single-cell transcriptomic atlas tracking the neural basis of division of labour in an ant superorganism. *Nat*. Ecol. Evol. 6, 1191–1204 (2022).

44. M. Neset Özel, F. Simon, S. Jafari, I. Holguera, Y.-C. C. Chen, N. Benhra, R. Naja El-Danaf, K. Kapuralin, J. Amy Malin, N. Konstantinides, C. Desplan, M. N. Özel, F. Simon, S. Jafari, I. Holguera, Y.-C. C. Chen, N. Benhra, R. N. El-Danaf, K. Kapuralin, J. A. Malin, N. Konstantinides, C. Desplan, Neuronal diversity and convergence in a visual system developmental atlas. Nature 589, 88–95 (2020).

45. M. Crow, A. Paul, S. Ballouz, Z. J. Huang, J. Gillis, Characterizing the replicability of cell types defined by single cell RNA-sequencing data using MetaNeighbor. Nat. Commun. 9, 884 (2018).

46. A. Borst, M. Helmstaedter, Common circuit design in fly and mammalian motion vision. Nat. Neurosci. 18, 1067–1076 (2015).

47. M. S. Maisak, J. Haag, G. Ammer, E. Serbe, M. Meier, A. Leonhardt, T. Schilling, A. Bahl, G. M. Rubin, A. Nern, B. J. Dickson, D. F. Reiff, E. Hopp, A. Borst, A directional tuning map of *Drosophila* elementary motion detectors. Nature 500, 212–216 (2013).

48. J. Erber, B. Pribbenow, A. Bauer, P. Kloppenburg, Antennal reflexes in the honeybee: tools for studying the nervous system. Apidologie 24, 283–296 (1993).

49. H. Kohno, S. Kamata, T. Kubo, Analysis of Antennal Responses to Motion Stimuli in the Honey Bee by Automated Tracking Using DeepLabCut. J. Insect Behav. 36, 332–346 (2023).

50. J. F. Nabhan, H. Pan, Q. Lu, Arrestin domain-containing protein 3 recruits the NEDD4 E3 ligase to mediate ubiquitination of the β2-adrenergic receptor. EMBO Rep. 11, 605–611 (2010).

51. M. Hammer, An identified neuron mediates the unconditioned stimulus in associative olfactory learning in honeybees. Nature 366, 59–63 (1993).

52. D. Owald, S. Waddell, Olfactory learning skews mushroom body output pathways to steer behavioral choice in Drosophila. Curr. Opin. Neurobiol. 35, 178–184 (2015).

53. R. L. Davis, Learning and memory using Drosophila melanogaster: a focus on advances made in the fifth decade of research. Genetics 224, iyad085 (2023).

54. J. Erber, P. Kloppenburg, The modulatory effects of serotonin and octopamine in the visual system of the honey bee (*Apis mellifera* L.) - I. Behavioral analysis of the motion-sensitive antennal reflex. J. Comp. Physiol. A 176, 111–118 (1995).

55. F. W. Schürmann, N. Klemm, Serotonin-immunoreactive neurons in the brain of the honeybee. J. Comp. Neurol. 225, 570–580 (1984).

56. I. Sinakevitch, M. Niwa, N. J. Strausfeld, Octopamine-like immunoreactivity in the honey bee and cockroach: Comparable organization in the brain and subesophageal ganglion. J. Comp. Neurol. 488, 233–254 (2005).

57. M. Thamm, S. Balfanz, R. Scheiner, A. Baumann, W. Blenau, Characterization of the 5-HT1A receptor of the honeybee (Apis mellifera) and involvement of serotonin in phototactic behavior. Cell. Mol. Life Sci. 67, 2467–2479 (2010).

58. M. V. Srinivasan, S. Zhang, M. Altwein, J. Tautz, Honeybee navigation: Nature and calibration of the “odometer.” Science 287, 851–853 (2000).

59. M. V. Srinivasan, S. Zhang, Visual motor computations in insects. Annu. Rev. Neurosci. 27, 679– 696 (2004).

60. J. Guo, A. Guo, Crossmodal interactions between olfactory and visual learning in *Drosophila*. Science 309, 307–310 (2005).

61. D. Thiagarajan, F. Eberl, D. Veit, B. S. Hansson, M. Knaden, S. Sachse, Aversive Bimodal Associations Differently Impact Visual and Olfactory Memory Performance in *Drosophila*. iScience 25, 105485 (2022).

62. A. Kamikouchi, H. Takeuchi, M. Sawata, S. Natori, T. Kubo, Concentrated expression of Ca2+/calmodulin-dependent protein kinase II and protein kinase C in the mushroom bodies of the brain of the honeybee *Apis mellifera* L. J. Comp. Neurol. 417, 501–510 (2000).

63. K. B. Gehring, K. Heufelder, I. Kersting, D. Eisenhardt, Abundance of phosphorylated *Apis mellifera* CREB in the honeybee’s mushroom body inner compact cells varies with age. J. Comp. Neurol. 524, 1165–1180 (2016).

64. T. Kuwabara, H. Kohno, M. Hatakeyama, T. Kubo, Evolutionary dynamics of mushroom body Kenyon cell types in hymenopteran brains from multifunctional type to functionally specialized types. Sci. Adv. 9, eadd4201 (2023).

65. H. Kohno, T. Kubo, mKast is dispensable for normal development and sexual maturation of the male European honeybee. Sci. Rep. 8, 11877 (2018).

66. H. Kohno, S. Suenami, H. Takeuchi, T. Sasaki, T. Kubo, Production of Knockout Mutants by CRISPR/Cas9 in the European Honeybee, *Apis mellifera* L. Zool. Sci. 33, 505–512 (2016).

67. S. W. Cobey, D. R. Tarpy, J. Woyke, Standard methods for instrumental insemination of *Apis mellifer*a queens. J. Apic. Res. 52, 1–18 (2013).

68. D. Conant, T. Hsiau, N. Rossi, J. Oki, T. Maures, K. Waite, J. Yang, S. Joshi, R. Kelso, K. Holden, B. L. Enzmann, R. Stoner, Inference of CRISPR Edits from Sanger Trace Data. Crisp. J. 5, 123– 130 (2022).

69. S. Suenami, S. Iino, T. Kubo, Pharmacologic inhibition of phospholipase C in the brain attenuates early memory formation in the honeybee (*Apis mellifera* L.). Biol. Open 7, bio028191 (2018).

70. S. Hori, H. Takeuchi, K. Arikawa, M. Kinoshita, N. Ichikawa, M. Sasaki, T. Kubo, Associative visual learning, color discrimination, and chromatic adaptation in the harnessed honeybee *Apis mellifera* L. J. Comp. Physiol. A Neuroethol. Sensory, Neural, Behav. Physiol. 192, 691–700 (2006).

71. A. Mathis, P. Mamidanna, K. M. Cury, T. Abe, V. N. Murthy, M. W. Mathis, M. Bethge, DeepLabCut: markerless pose estimation of user-defined body parts with deep learning. Nat. Neurosci. 21, 1281–1289 (2018).

72. A. M. Bolger, M. Lohse, B. Usadel, Trimmomatic: a flexible trimmer for Illumina sequence data. Bioinformatics 30, 2114–2120 (2014).

73. D. Kim, B. Langmead, S. L. Salzberg, HISAT: a fast spliced aligner with low memory requirements. Nat. Methods 12, 357–360 (2015).

74. M. Pertea, G. M. Pertea, C. M. Antonescu, T. C. Chang, J. T. Mendell, S. L. Salzberg, StringTie enables improved reconstruction of a transcriptome from RNA-seq reads. Nat. Biotechnol. 33, 290– 295 (2015).

75. J. Sun, T. Nishiyama, K. Shimizu, K. Kadota, TCC: An R package for comparing tag count data with robust normalization strategies. BMC Bioinformatics 14, 1–14 (2013).

76. Y. Zhou, B. Zhou, L. Pache, M. Chang, A. H. Khodabakhshi, O. Tanaseichuk, C. Benner, S. K. Chanda, Metascape provides a biologist-oriented resource for the analysis of systems-level datasets. Nat. Commun. 10, 1523 (2019).

77. K. Emoto, S. Yamashita, Y. Okada, Mechanisms of heat-induced antigen retrieval: Does pH or ionic strength of the solution play a role for refolding antigens? J. Histochem. Cytochem. 53, 1311– 1321 (2005).

78. B. R. E. Klagges, G. Heimbeck, T. A. Godenschwege, A. Hofbauer, G. O. Pflugfelder, R. Reifegerste, D. Reisch, M. Schaupp, S. Buchner, E. Buchner, Invertebrate Synapsins: A Single Gene Codes for Several Isoforms in Drosophila. J. Neurosci. 16, 3154–3165 (1996).

79. R. Satija, J. A. Farrell, D. Gennert, A. F. Schier, A. Regev, Spatial reconstruction of single-cell gene expression data. Nat. Biotechnol. 33, 495–502 (2015).

80. P. Shannon, A. Markiel, O. Ozier, N. S. Baliga, J. T. Wang, D. Ramage, N. Amin, B. Schwikowski, T. Ideker, Cytoscape: A Software Environment for Integrated Models of Biomolecular Interaction Networks. Genome Res. 13, 2498–2504 (2003).

